# Modulatory mechanisms of TARP γ8-selective AMPA receptor therapeutics

**DOI:** 10.1101/2022.09.25.509219

**Authors:** Danyang Zhang, Remigijus Lape, Saher A. Shaikh, Bianka K. Kohegyi, Jake F. Watson, Ondrej Cais, Terunaga Nakagawa, Ingo H. Greger

**Affiliations:** Neurobiology Division, MRC Laboratory of Molecular Biology, Cambridge, UK; IST Austria, Klosterneuburg, Austria; Department of Molecular Physiology and Biophysics, Vanderbilt University, School of Medicine, Nashville, USA

**Author notes:** These authors contributed equally (Danyang Zhang, Remigijus Lape).

## Abstract

AMPA glutamate receptors (AMPARs) mediate excitatory neurotransmission throughout the brain. Their signalling is uniquely diversified by brain region-specific auxiliary subunits, providing an opportunity for the development of selective therapeutics. AMPARs associated with TARP γ8 are enriched in the hippocampus, and are targets of emerging anti-epileptic drugs. To understand their therapeutic activity, we determined cryo-EM structures of the GluA1/2-γ8 receptor associated with three potent, chemically diverse drugs. We find that despite sharing a lipid-exposed and water-accessible binding pocket, drug action is differentially affected by binding-site mutants. Together with patch-clamp recordings and MD simulations we demonstrate that ligand-triggered reorganisation of the AMPAR-TARP interface contributes to modulation. Unexpectedly, one ligand (JNJ-61432059) acts bifunctionally, negatively affecting GluA1 but exerting positive modulatory action on GluA2-containing AMPARs, in a TARP stoichiometry-dependent manner. These results further illuminate the action of TARPs, demonstrate the sensitive balance between positive and negative modulatory action, and provide a mechanistic platform for development of both positive and negative selective AMPAR modulators.

## Introduction

AMPARs mediate the majority excitatory synaptic transmission in the brain, and are central to the synaptic plasticity that underlies learning ^1^. Their dysfunction (both hypo- and hyper-excitability) is associated with multiple neurological and neuropsychiatric disorders, including memory deficits, epilepsy, depression, motoneuron disease and gliomas ^1-3^. Due to their unique molecular diversity, AMPARs generate a rich repertoire of post-synaptic response properties ^1,4^, consequently offering sites for selective therapeutic intervention ^5-9^. Four core subunits (GluA1-4) arrange as two non-equivalent pairs (termed AC and BD) that fulfil different functions ^10,11^. Further diversity comes from a wide array of auxiliary subunits, associating with the receptor in various stoichiometries, and in a circuitry-dependent fashion ^4,12-15^.

Major auxiliary subunits associate with the ion channel periphery, including the TARPs (transmembrane AMPAR regulatory proteins), CNIHs (cornichon homologs), and GSG1L (germline-specific gene 1) ^1,16,17^. TARPs are the most widely expressed, and constitute an integral component of AMPARs. Resembling tetraspan claudins^18^, TARPs have four transmembrane helices and an extracellular, five-stranded beta-sheet harbouring flexible loops ^10,19,20^. Based on protein sequence and function, three groups exist, Type 1a (TARPs γ2, γ3), Type 1b (γ4, γ8), and Type 2 (γ5, γ7), which are expressed in overlapping yet distinct brain regions, differentially impacting gating kinetics, ion conductance, pharmacology, and trafficking ^4,12,13,16,21^.

A maximum of four TARPs associate with the receptor through two pairs of binding sites (termed A’C’ and B’D’) (**Fig. 1**), formed by the gate-surrounding M1 and M4 helices ^19,20^. The more accessible B’D’ sites are preferentially occupied by the bulkier TARPs (Type 1b and Type 2) ^22,23^, and by GSG1L ^24^, while Type 1a TARPs and CNIHs have no obvious site preference ^19,25^. TARP stoichiometry determines receptor function, as the twofold symmetry of the receptor dictates that the two pairs of binding sites provide different access for the TARP to the gating machinery ^19,22^, consisting of the M3 gating helices, the gating linkers, and the ligand-binding domains (LBDs). TARP interactions with the receptor are complex: the TARP M3 and M4 helices engage the ion channel through transmembrane contacts, while their extracellular loops mediate transient contacts with the LBD and the gating linkers ^1,16,26^. How this arrangement generates the wide spectrum of TARP modulation remains to be resolved.

**Fig. 1:**
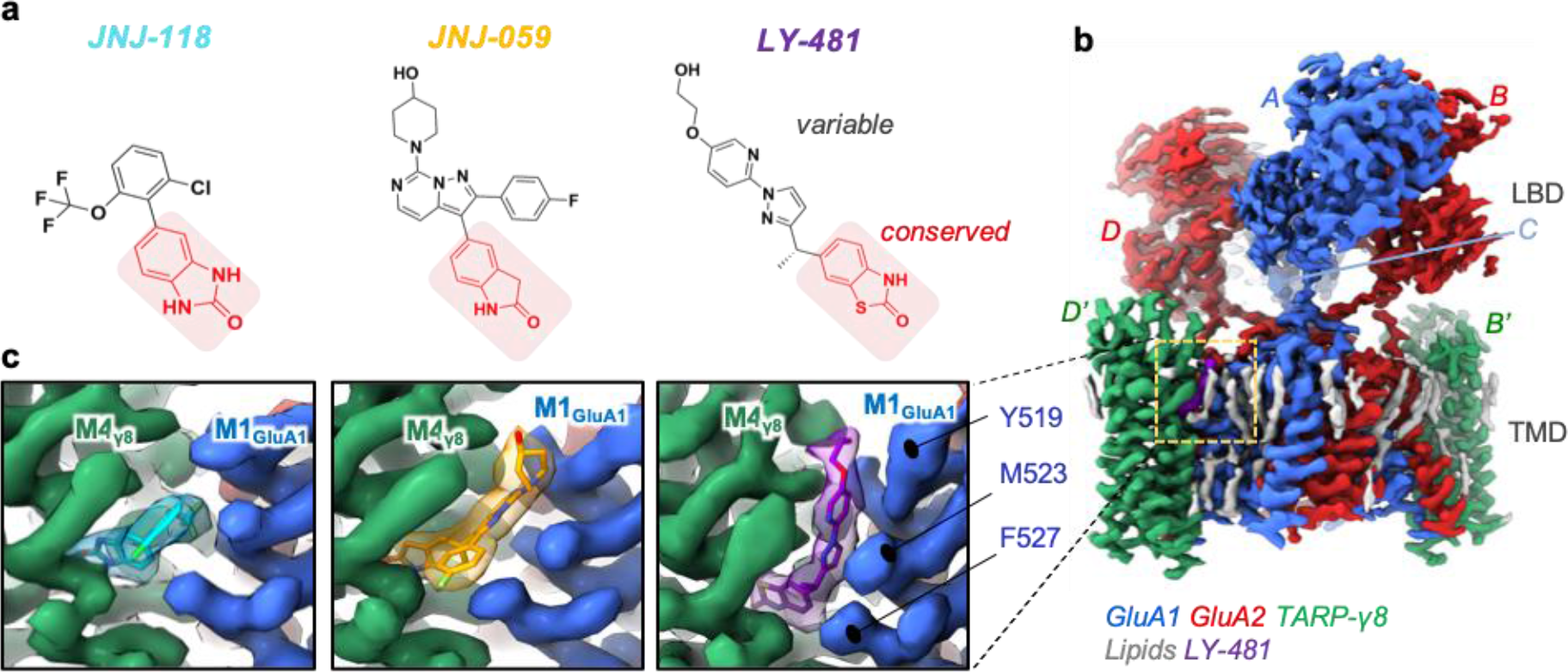
Cryo-EM maps of three NAMs bound to the resting-state GluA1/2_γ8 AMPAR. **a**, Chemical structures of the three NAMs. The shared oxindole isostere is labelled red, the variable groups are shown in black. **b**, Cryo-EM map of the GluA1/2_γ8 LBD (Ligand Binding Domain) and TMD (Transmembrane Domain) sectors, associated with LY-481. The colour code of protein, lipid and NAM is indicated below the map. Subunits are also labelled as described in the text (GluA1: A, C; GluA2: B, D; TARP γ8: B’, D’). **c**, Cryo-EM maps of each NAM in side view. TARP γ8 (helix M4) is shown in green, GluA1 (helix M1) in blue. Also shown are three GluA1 residues involved in NAM co-ordination (Tyr519, Met523, Phe527).

As major excitatory receptors, AMPARs are central drug targets. However, the majority of available drugs target the core subunits, thus causing severe side effects due to their ubiquitous presence across the brain. These include negative allosteric modulators (NAMs) such as the anti-epileptic perampanel ^27,28^, while positive allosteric modulators (PAMs) are also being developed as cognitive enhancers ^29,30^. By contrast, auxiliary subunit diversity offers scope for the development of region-selective AMPAR therapeutics. Recently discovered NAMs target γ8, a TARP strongly enriched in the hippocampus, and show promise in pre-clinical epilepsy studies ^31,32^. One example, LY-3130481 (CERC-611), is also effective in the treatment of pain and of gliomas ^33,34^, with Phase-2 clinical trials currently ongoing (https://clinicaltrials.gov/ct2/show/NCT04714996). Understanding the binding and mechanism of these drugs, will not only facilitate the refinement and development of novel therapeutics, but also enables unlocking the complexity of TARP action.

Here, high-resolution cryo-EM structures of GluA1/2 TARP-γ8 AMPAR complex, associated with three γ8 drugs, differing in size and potency: LY-3130<u>481</u>, JNJ-55511<u>118</u>, JNJ-61432<u>059</u> (herein referred to as ‘LY-481’, ‘JNJ-118’, ‘JNJ-059’), reveal how these NAMs engage the channel, and reshape the interface between the AMPAR and γ8. Structures of active and resting states, combined with all-atom molecular dynamics (MD) simulations and electrophysiological studies provide a basis of NAM action, and reveal a surprising receptor- and ligand-specific PAM activity, operating through a different allosteric route. In addition to further advancing our understanding of AMPAR modulation by γ8, our results have broader implications for the development of highly specific AMPAR therapeutics.

## Results

LY-481 and JNJ-118 were identified in high-throughput screens, while JNJ-059, a substantially more potent ligand, was developed subsequently (**Fig. 1a**) ^31,32,35,36^. All three compounds have a shared oxindole group, which appears fundamental to TARP γ8 binding in our previous modelling study ^37^. To understand and compare the binding modalities of these compounds, we obtained cryo-EM structures of each NAM with resting-state GluA1/2 receptors (**Table S1**), containing two γ8 subunits at the preferred B’D’ sites (Methods) (**Fig. 1b**). JNJ-059 was also captured in an active state, with an open channel gate. Resolutions of the ion channel/TARP sector at **∼** 3.0 Å, with the LY-481 structure reaching 2.6 Å, enabled an unprecedented view of the modulator binding site with associated lipids and putative water molecules (**Fig. S1-3**, and **Supplemental Movie 1**).

In addition, we subjected these structures to large scale MD simulations in an explicit lipid membrane together with water and ions, alongside previously reported apo-state complexes (open and resting; PDB: 7QHB and 7OCD, respectively) ^22,38^. Three sets of simulations for each receptor, totaling 1.25-1.5 μs of sampling for each system, offered additional insight into ligand behavior in their binding pocket, as well as enabling a comprehensive comparison of local and global receptor dynamics in response to NAM binding (**Supplemental Movie 2**).

### Ligand binding mode

The NAM binding pocket locates near the boundary of the lipid bilayer with the extracellular milieu (**Figure 1b** and **c**). MD simulations reveal water molecules penetrating into the pocket from γ8 Ser128 on the extracellular edge down to γ8 Asn172 in the binding pocket (**Fig. S4a**), suggesting a potential route for ligand entry. Putative waters are also apparent in the LY-481 cryoEM map around GluA2 Ser790, and in the NAM pocket at γ8 Ser128 (**Fig. S4b and c**).

Modulator selectivity for TARP γ8 is determined by two residues unique to γ8, Val176 and Gly209, in the M3 and M4 helices ^31,32^. These positions are occupied by bulkier residues in the other TARPs (Ile and Ala, respectively), restricting ligand access (**Fig. S4d**). All three NAMs anchor between these two γ8 residues through a common oxindole isostere secured by a H-bond between γ8 Asn172 and an amine of the oxindole (**Fig. 2, and Fig. S4e**), consistent with our recent docking study ^37^, and replicating a previously reported binding mode of JNJ-118 ^39^. The oxindole’s benzene ring is capped by γ8 Phe205, further stabilizing the ligand. MD simulations show that the ligands remain stably bound in this site throughout 500 ns runs (**Fig. S4f**).

**Fig. 2:**
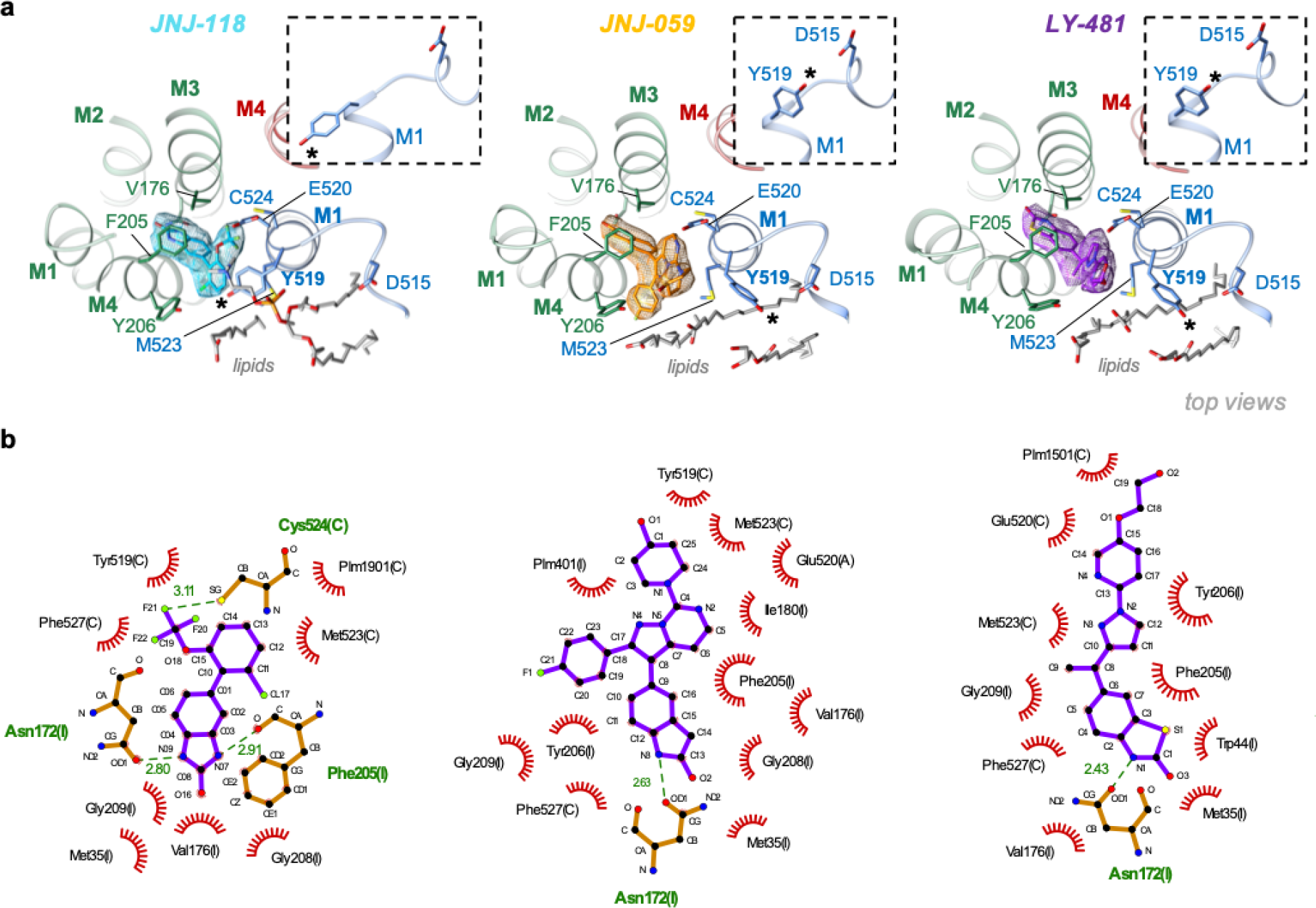
NAM binding pockets and coordinating residues. **a**, Top view onto the NAM pockets. TARP γ8 is shown in green, AMPAR helices are blue (GluA1 M1) and red (GluA2 M4), the three NAMs are colour-coded. Ligand-interacting residues and lipids are shown in stick. The insets show side views onto Tyr519, which swings towards Asp515 in the JNJ- 059 and LY-481 structures (Tyr519 are also highlighted with asterisks). **b**, Liplot diagrams of the three NAMs ^55^. H-bonds are indicated as green, dashed lines, while non-bonded contacts with the ligands are denoted by red spoked arcs.

While oxindole groups are shared, and their binding modes analogous, the remainder of each compound is variable. These variable moieties project towards the top of the GluA1 M1 helix, increasing the distance between M1 and the γ8 M4 helix, and triggering side chain reorientations of Tyr519, Met523 and Phe527 (**Fig. 2a**). The same residues rearrange to engage γ8 upon binding to the receptor ^10^. Together with Glu520 and Cys524, they form the main ligand coordination points on the AMPAR (**Fig. 2b**). In the JNJ-118 structure, the top of the binding site is capped by Tyr519, which points towards γ8 Tyr206, comparable to ligand-free structures. By contrast, the two larger NAMs reach towards Glu520 at the lipid-solvent interface, and force the Tyr519 side chain towards Asp515 in the pre-M1 helix (**Fig. 2a**). Pre-M1 surrounds the M3 gating helices ^11^, and dilates together with the M3 gate on channel opening ^22^, thus forming a key regulatory element adjacent to the NAM pocket (**Fig. 3a**). Tyr519 persistently engages Asp515 through a water-mediated H-bond in MD runs with JNJ-059 or LY-481 bound, but this interaction is not observed with either a ligand-free structure (PDB: 7OCD), or with the smaller JNJ-118 ligand, where Tyr519 engages γ8 Tyr206 instead (**Fig. 3a-c**). Notably, a putative H-bond between Tyr519 and Asp515 is also apparent in our high-resolution LY-481 structure (**Fig. S5a**). We hypothesize that this bond impacts the local dynamics between pre-M1 and the M3 gate, perhaps by influencing a hydrophobic interaction between the neighboring Pro516 (in pre-M1) and Phe619 (in M3). These two highly conserved residues are implicated in gating regulation in NMDA receptors ^40,41^, and in coordinating the negative allosteric AMPAR modulators GYKI and perampanel, binding the core subunits ^42,43^.

**Fig. 3:**
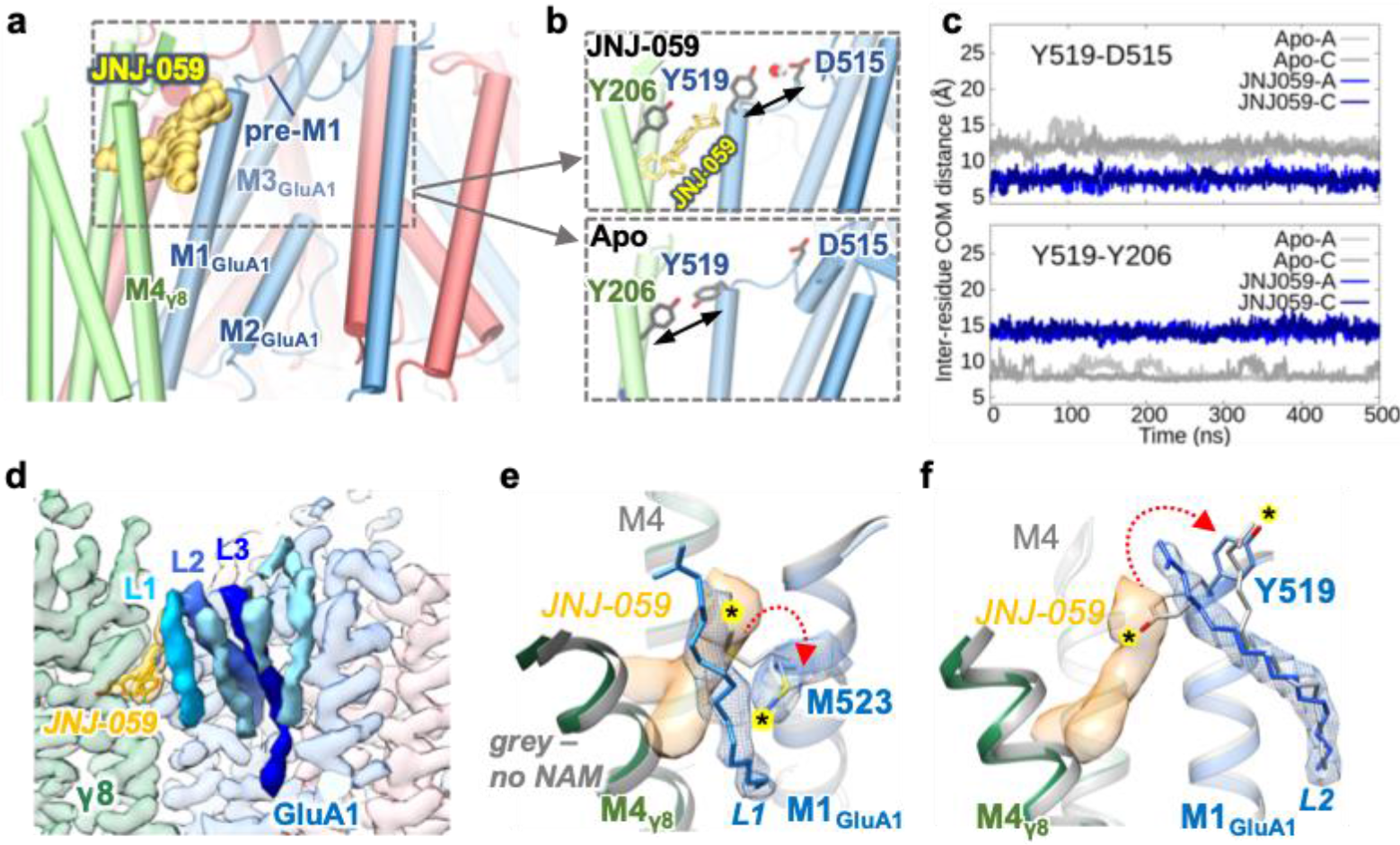
JNJ-059-induced H-bond formation, and lipid interactions. **a**, Model of the receptor complex with JNJ-059 (yellow spheres), highlighting the proximity between the NAM pocket and the GluA1 pre-M1 helix (the figure is derived from an MD simulation). **b**, JNJ-059 disrupts Tyr519 interaction with Tyr206 (γ8) (top panel), as compared to a simulation without the NAM (‘Apo’; bottom panel), but enables an H-bond between Tyr519 and Asp515 in the pre-M1 helix (top). **c**, Variations in distance between centres of mass for Tyr519-Asp515 and Tyr519-Tyr206 in MD simulations of resting-state models. Distances are compared for the apo state (grey) and the JNJ059- bound structure (blue). Data are for both GluA1 subunits (chains A and C) shown in light and dark colours. **d**, Cryo-EM map of the resting-state JNJ-059 structure. Lipid densities stacking along the GluA1 pre- M1 helix are shown in shades of blue (L1-3). **e**,**f**, Overlay between the JNJ-059-resting model (coloured) with an apo resting state (PDB: 7OCD; grey). The NAM induces an interaction between Met523 and the acyl chain of L1 (E), and a rearrangement of L2 via the Tyr519 side chain (F). NAM-induced side chain reorientations are highlighted with an asterisk. Note the dilation of the NAM pocket in the presence of JNJ-059.

### Impact of JNJ-059 on opening of the gate

How these NAMs affect opening of the gate is currently elusive. To address this, we determined an open-state structure of GluA1/2_γ8 associated with JNJ-059, which was overall similar to an open-state without the modulator (PDB: 7QHB); with a root mean square deviation of 0.36 Å, when aligning Cα atoms of the AMPAR/TARP transmembrane sector (**Fig. S5b**). When focussing onto the gating core, opening of the M3 gate was still apparent in the presence of JNJ-059, although the gate constriction point (at GluA1 Thr621/GluA2 Thr625) was slightly wider without the NAM (by ∼0.5 Å) (**Fig. S5c** and **d**). Therefore, even the bulkiest NAM does not severely compromise an open gate conformation, underscoring mechanistic differences to NAMs targeting the core receptor such as perampanel ^43,44^. Based on a resting state structure in complex with JNJ-118, it has been hypothesized that NAM binding precludes outward movement of the M3 gating helices ^39^. However, the highly similar architecture of active states in the absence and presence of a NAM indicates other contributors underlying NAM action, as investigated below.

### Lipids shape the NAM-binding pocket

The binding pocket opens sideways into the lipid bilayer, enabling ligand interactions with lipids, which stack nearby along the horizontal pre-M1 helices (**Fig. 3d**) ^10^. The fluorophenyl group of JNJ-059 projects towards the acyl chain of a lipid (L1) (**Fig. 3e**), which is also engaged by the distal hydroxy-ethyloxy portion of LY-481, close to its (currently unresolved) head-group (**Fig. S5e**). In the JNJ-118 structure, L1 arches above Tyr519, effectively capping the binding pocket (**Fig. S5f**). L1 is also contacted by the displaced Met523 side chain in all three NAM structures, while the neighbouring lipid (L2) reorients around the dynamic Tyr519 side in response to LY-481 and JNJ-059 (**Figure 3e, f and Fig. S5e**). Hence, the three NAMs perturb the local lipid environment with the AMPAR in different ways. As these lipids (L1-L3) link the pre-M1 helix to the helices lining the conduction path - the M2 pore helix of the selectivity filter and M3 gating helix (**Fig. S5g**) ^22^, ligand-induced lipid reorganisation is a possible contributer to NAM action.

### Interfering with NAM action

To assess AMPAR modulation by the three NAMs comparatively we used patch-clamp recordings of recombinantly expressed receptor complexes. First, we generated a baseline for the action of γ8, using high-resolution recordings of GluA1 fused to γ8 (denoted GluA1_γ8) (**Fig. S6a**). Since kinetics and drug effects were similar when applying glutamate to either isolated membrane patches or to whole HEK293 cells (**Fig. S6a**,**b**), we used the latter approach throughout this study. As expected, all three NAMs (at 10 μM) reduced the positive modulation conferred by γ8: reducing peak current amplitude, quickening desensitization kinetics, and decreasing equilibrium current and resensitization. All together, these changes reduce charge transfer through the channel (**Fig. 4a**, and **Fig. S6c**). Despite their structural differences, the extent of NAM action was similar for all three NAMs.

**Fig. 4:**
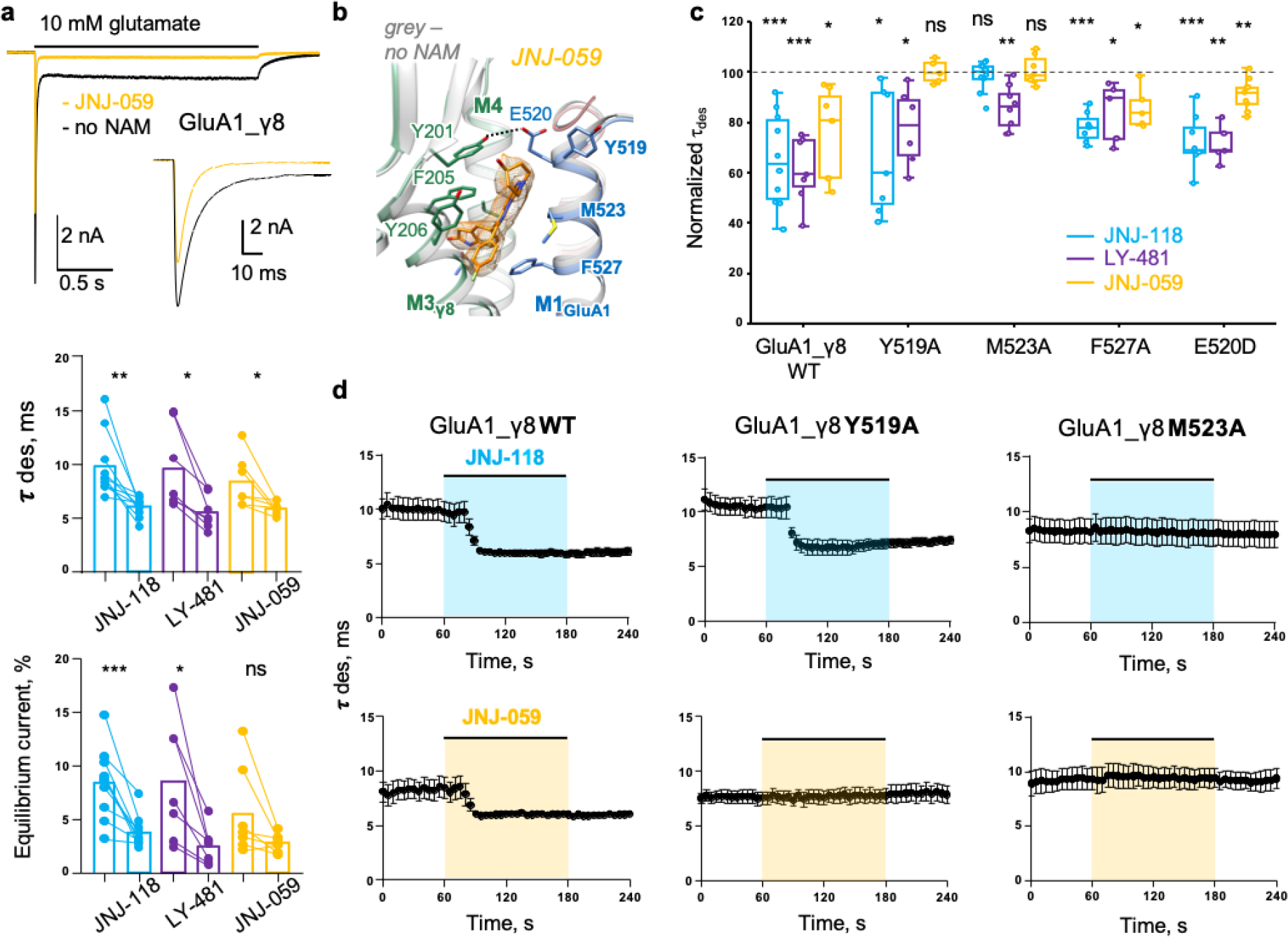
Functional NAM effects on wild type and mutant GluA1_γ8. **a**, Whole-cell response of GluA1_γ8 to 10 mM glutamate (2 sec; -60 mV), in either the absence (black) or presence (yellow) of JNJ-059. The inset zooms into the peak response. Middle and bottom: Paired plots showing effect of NAMs on desensitization τ (Middle; JNJ-118: control 9.8±0.9 ms, JNJ- 118 6.0±0.3 ms, n=10; LY-481: control 9.6±1.5 ms, LY-481 5.6±0.6 ms, n=7; JNJ-059: control 8.4±0.9 ms, JNJ-059 6.0±0.2 ms, n=7), and the equilibrium current (Bottom; JNJ-118: control 8.4±1.0%, JNJ- 118 3.8±0.5 ms, n=10; LY-481: control 8.5±2.1 ms, LY-481 2.5±0.7 ms, n=7; JNJ-059: control 5.6±1.6 ms, JNJ-059 2.7±0.3 ms, n=7). Black circles show individual values, bar height indicates the mean value. Asterisks indicate summary of two-tailed paired t-test values: * *p* <= 0.05, ** *p* <= 0.01, *** *p* <= 0.001 and ‘ns’ for p>0.05. **b**, JNJ-059-induced rearrangement of residues along the GluA1 M1 helix. A ligand-free apo structure is superimposed (grey). The stippled line indicates a potential H-bond between Tyr201 and Tyr519. **c**, Summary data for GluA1_γ8 wild-type and M1 mutants Box plots showing extent of macroscopic desensitization τ change after application of JNJ-118, LY-481 or JNJ-059 for wild type or mutant GluA1_γ8. Each point is a desensitization τ measured in presence of modulator normalized to the pre-application value and expressed as percentage (WT: JNJ-118 65±6% n=10, LY-481 61±5% n=7, JNJ-059 75±7% n=7; Y519A: JNJ-118 68±9% n=7, LY-481 78±6% n=6, JNJ-059 100±2% n=5; E520D: JNJ-118 72±4% n=8, LY-481 72±3% n=5, JNJ-059 91±2% n=9; M523A: JNJ-118 99±2% n=10, LY-481 86±3% n=8, JNJ-059 101±2% n=7; F527A: JNJ-118 78±2% n=7, LY-481 84±5% n=5, JNJ-059 86±4% n=5). Boxes show the 25^th^/75^th^ percentiles and whiskers indicate the furthest points that fall within 1.5 times of interquartile range from the 25^th^/75^th^ percentiles. The horizontal line in each box shows the median value. Asterisks summarize one-sample t-test (difference from 100%) results (see (a) for details). **d**, Time course of desensitization entry in response to JNJ-118 (top row) and JNJ-059 (bottom) row for GluA1_γ8 (left), Y519A mutant (middle) and M523 mutant (right). Scatter plot of average desensitization τ in time indicating the time-course of JNJ-059 or JNJ-118 effect on wild-type, M523A or Y519A GluA1i_ γ8. Black circles and whiskers indicate average values and standard error. Number of cells are n=10 for GluA1i_γ8, n=7 for GluA1i _γ8 Y519A, n=10 for GluA1i_γ8 M523A with JNJ-118 and n=7 for GluA1i_γ8, n=5 for GluA1i_γ8 Y519A, n=7 for GluA1i_γ8 M523A with JNJ-059.

Differences became apparent when mutating side chains in the M1 helix that are displaced by the NAMs (**Fig. 4b, c**, and **Fig. S6d**). At the top of M1, mutation of the conformationally variable Tyr519 to alanine (Y519A) (**Fig. 2a** and **3b**) abolished negative modulation (desensitization kinetics and the equilibrium response) of JNJ-059 (100 ± 2%, n=5, *p*=0.9 one-sample two-tail t-test comparison to 100%), but not of JNJ-118 (68 ± 9%, n=7, *p*=0.01) and LY-481 (78 ± 6%, n=6, *p*=0.02) (**Fig. 4c, d and Fig. S6d**), suggesting that ligand-mediated projection of the Tyr519 side chain toward the pre-M1 helix is critical for JNJ-059 action. Mutation of Met523 (M523A) essentially blunted NAM action of both JNJ-118 and JNJ-059 but not of LY-481 (**Fig. 4c** and **d**). All NAMs break the γ8 Phe205 GluA1 Met523 interaction and force the Met523 side chain into lipid (**Fig. 3e**). The lipid interactions induced may contribute to negative modulation, and are lost by mutation to the shorter alanine side chain. F527A was of lesser impact across the three NAMs. Together, modulatory action is sensitive to features of the NAM variable groups, and their interaction pattern with the M1 helix, leading to ligand-specific local signalling routes (**Fig. S6d**). This is highlighted further by the E520D mutation, which ruptures an H-bond between Glu520 and γ8 Tyr201, and selectively impacts the larger LY-481 and JNJ-059 NAMs (**Fig. 4b and c**).

### Endogeous oxindoles target the NAM pocket

To further investigate the specific modulatory properties of the NAM variable groups, we studied the effect of oxindole derivatives lacking this extension (**Fig. S7a**). The naturally occurring tryptophan metabolites isatin and 5-OH-indole are structurally highly similar to the γ8 docking group of the NAMs. These compounds also docked between the γ8 M3 and M4 helices (the cognate NAM site) in MD simulations, and remained bound throughout the run (**Fig. S7b**). Interestingly however, we observed no functional effect of their binding in patch-clamp recordings with GluA1_γ8 (**Fig. S7a**). Therefore, simply binding of the oxindole to the TARP is not sufficient for NAM action - projection of the variable group towards the receptor is essential for negative modulation by these compounds.

### Modulatory role of the γ8 transmembrane sector

The structural changes upon NAM binding are not limited to the side chain level. We observe global changes in the arrangement between the receptor and the TARP, which likely contribute to negative modulation. Specifically, a pronounced vertical outward rotation of γ8 helices in NAM-bound structures, relative to apo structures, is apparent (**Fig. 5a**). This motion was most prominent with JNJ-059, and includes the peripheral M1 and M2 helices as well as the top of M4, but not M3, which likely acts as a pivot axis. The clockwise rotation (when viewed from the top) could counter the motion of the TARP towards the receptor that accompanies channel activation ^22^, and may thereby compromise transition to the active state, and/or its stabilization.

**Fig. 5:**
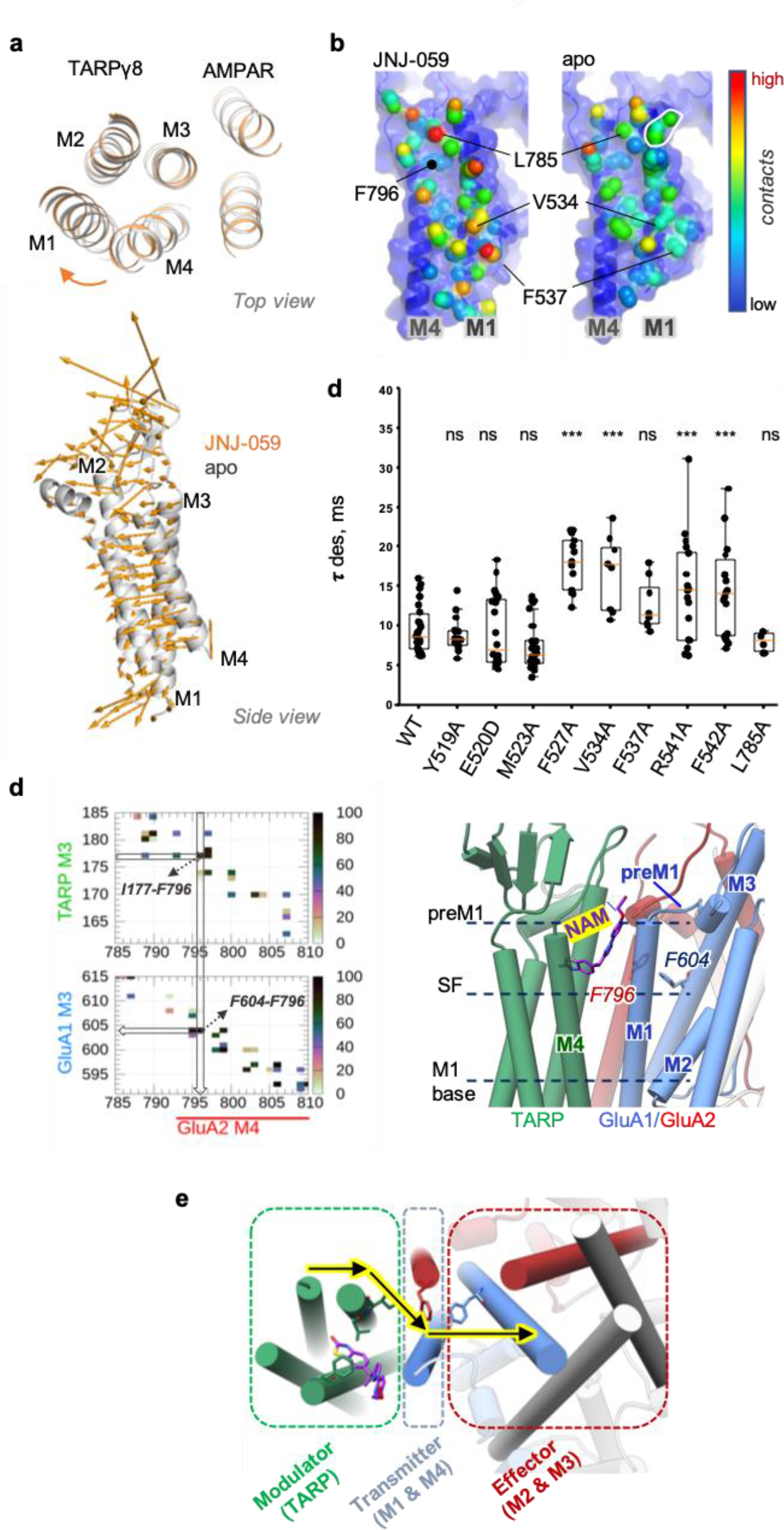
NAMs trigger a global reorientation of TARP γ8. **a**, JNJ-059-induced rotation of γ8 helices M1, M2 and M4, relative to an apo structure (PDB: 7OCD). Alignment of the the TMD sector of apo (grey) and JNJ-059-bound (orange) resting-state models. The vectors indicate the direction of γ8 in response to the NAM in both top and side views. Vectors were generated using the ‘modevectors’ script in PyMol. **b**, TARP γ8 contact points along its binding site, the M4_GluA2_ and M1_GluA1_ helices. Contacted residues are coloured depending on the number of atoms contributing to the interaction (red: high; blue: low). Countacts were computed using ‘findNeighbors’ in ProDy’ with a 4.5 Å cutoff between heavy atoms ^56^. **c**, Box plots showing macroscopic desensitization τ for GluA1_γ8 wild type and mutants; each point is a desensitization τ (WT: 9.7±0.6 ms, n=30; Y519A: 8.9±0.5 ms, n=18; E520D: 9.4±1.0 ms, n=22; M523A: 7.4±0.6 ms, n=27; F527A: 18.0±0.9 ms, n=13; V534A: 17±2 ms, n=8; F537A: 13±1 ms, n=8; R541A: 16±1 ms, n=24; F542A: 15±1 ms, n=21; L785A: 8.0±0.5 ms, n=6) obtained by fitting the decaying phase of whole-cell currents with a single exponential. Boxplots are as described in Fig 4C. Asterisks summarize one-way ANOVA test, Dunnett correction was used for multiple comparisons to wild type receptor (see Fig. 4a for details). **d**, High-frequency residue contacts forming a potential pathway from the NAM binding site to the gate. Left panel: TARP-GluA2 (top) and GluA2-GluA1 (bottom) contact maps for the JNJ-059 resting state MD simulation, suggesting a route from TARP Ile177 via GluA2 Phe796 to GluA1 (see Methods for further detail). Middle panel: Pathway residues relative to key regions in the AMPAR pre-M1, selectivity filter (SF), and cytosolic base of M1. **e**, Overall model suggesting that allosteric information from the TARP (Modulator) is communicated via the AMPAR peripheral M1+M4 helices (Transmitter) to the M3 gate (Effector).

The three NAMs also alter the interaction landscape between γ8 and its binding site, the GluA1 M1 and GluA2 M4 helices. Contrary to what could be expected, contact analyses reveal *enhanced* interaction points with the receptor in the NAM structures, rather than a decoupling of γ8 (in comparison to apo structures PDB: 7OCD and 7OCE) (Methods) (**Fig. 5b** and **Fig. S7c**). Strengthened contacts include Phe527, Val534 and Phe537 on the M1 helix, as well as Leu789, Val800 and Met807 on M4. Side chain interactions immediately adjacent to the binding pocket are either lost (Tyr519, Met523) or are unaltered (Cys524). This NAM-triggered realignment of the TARP-AMPAR complex likely alters the overall energetics of the system.

To extend these findings, we examined AMPAR-TARP interaction dynamics through all-atom MD simulations. We computed AMPAR-TARP contacts, and generated difference maps between simulations from NAM-bound versus an apo structure (PDB: 7OCD) (Methods). This analysis further illuminated the NAMs-induced rearrangement of the complex; while contacts around the NAM pocket are lost (GluA1 Tyr519, Glu520, Met523), the base of GluA1 M1, around Phe542, forms more persistent interactions with the γ8 M4 helix. Meanwhile, the γ8 M3 helix improves its overall engagement with GluA2 M4 at Leu789/Ser790, Val800, Met807 (**Fig. S7d**). The enhanced contacts at the base of GluA1 M1 with the TARP extend into the cytoplasmic loop connecting M1 with the M2 pore helix, which forms the selectivity filter. Phe542, and the neighbouring Arg541, re-orient their side chains on γ8 binding ^22^, with Arg541 poised to engage the negatively charged M1/M2 loop. These two M1 residues have a substantial impact on γ8 modulation, as we show below.

To address whether rearrangement of γ8 contributes to modulation, we mutated multiple GluA1 residues, scattered throughout the binding-site, and recorded their functional effects in the absence of NAMs. This analysis revealed bimodal effects on γ8 function: while M523A, E520D and the nearby L785A tended to reduce positive modulation by the TARP, other mutants further slowed desensitization kinetics and increased the equilibrium response (**Fig. 5c** and **Fig. S8a**). Positive modulation is facilitated by F527A and by mutants in the lower part of M1, including V534A, F537A, R541A and F542A. Interestingly, this pattern correlates with M1-TARP interaction strength – the upper part of M1 is loosely coupled to γ8, while the region below Phe527 is coupled more tightly ^38^. Moreover, this enhanced TARP modulation is not immune to NAM action, as positions distal from the binding pocket are also robustly blunted by all three NAMs, as shown for R541A and F542A (**Fig. S8b**). Therefore, in addition to the more intensively studied TARP loops, our analysis highlight transmembrane interactions in TARP modulation, which are sensitive to perturbation of individual contact points.

### Mechanism of NAM action

Taken together, we propose that γ8 NAMs signal through a combination of local and global routes. Due to their proximity to the gate, local perturbations can be transmitted effectively, via the top of M1 and the pre-M1 helix (**Fig. 3a**). These perturbations include side chains of binding pocket residues, such as Met523 and Tyr519, as well as associated lipids stacking along pre-M1 (**Fig. 2a** and **3d-f**). This gate-proximal region is engaged differently by other major AMPAR auxiliary subunits, such as CNIHs and GSG1L. By influencing either transition to the active state, or the stability of the open state conformation ^22^, they generate their unique modulatory profiles ^22,25,45,46^. NAMs target this strategic location and interfere with this process.

In addition, the NAM-triggered realignment of the TARP can be communicated across the entire γ8 binding-site (the M1, M4 helices) to both elements of the conduction path, the M3 gate and the M2 pore loop. Possible transmission pathways between contact points emerge from MD simulations, where residues contributing to a given pathway are identified through stable interaction times during the MD simulations (i.e. high percentage of residue pair interactions within 4Å, excluding hydrogens) (**Fig. 5d;** left panel). One observed route leads from the ligand co-ordinating Val176/Ile177 in γ8, via Phe796 (in the GluA2 M4 helix) to Phe604 in the M3 helix (**Fig. 5d**). Phe796 is strategically located within van der Waals contact distance of both the TARP and the M3 helix (at Phe604), as well as the NAM coordinating Cys524 (**Fig. 2c** and **5b**). Towards the cytoplasmic end of M1, residues sensitive to modulation by γ8 (such as Phe537; **Fig. 5c**) directly contact the M2 pore loop of the selectivity filter and will thereby influence the lower half of the conduction path (**Fig. 5d;** right panel), while Arg541 targets the acidic M1/2 cytoplasmic loop, implicated in AMPAR regulation ^38,47^. In summary, contacts in the AMPAR-TARP transmembrane sector appear to be finely tuned for transmission via the M1/M4 periphery (‘transmitter’) to the ‘effector’ (the M3 gate and M2 selectivity filter) (**Fig. 5e**). This interaction landscape is perturbed by NAM binding.

### JNJ-059 acts as a PAM on GluA2

We obtained an unexpected result when recording the action of JNJ-059 on a GluA2 γ8 homomer (GluA2 fused to γ8; GluA2_γ8), which was *opposite* to what was seen with GuA1_γ8. Contrary to the speeding of desensitization, which is expected by interference with γ8’s modulatory action, JNJ-059 triggered a slowly developing PAM response with GluA2_γ8, manifested in further slowing desensitization kinetics (**Fig. 6a**). JNJ-059 also failed to reduce the GluA2_γ8 equilibrium current, but still abolished resensitization (**Fig. S9a**). This behaviour was not seen with JNJ-118, which acted as a NAM on GluA2 for all measures (**Fig. S9a** and **b**), consistent with a recent study ^48^. Similar to JNJ-118, LY-481 triggered a pronounced reduction of the equilibrium response, but modestly slowed GluA2_γ8 desensitization kinetics (**Fig. 6b**). Hence, the two larger ligands failed to exert a full NAM effect on GluA2, with JNJ-059 binding at this site also generating PAM activity.

**Fig. 6:**
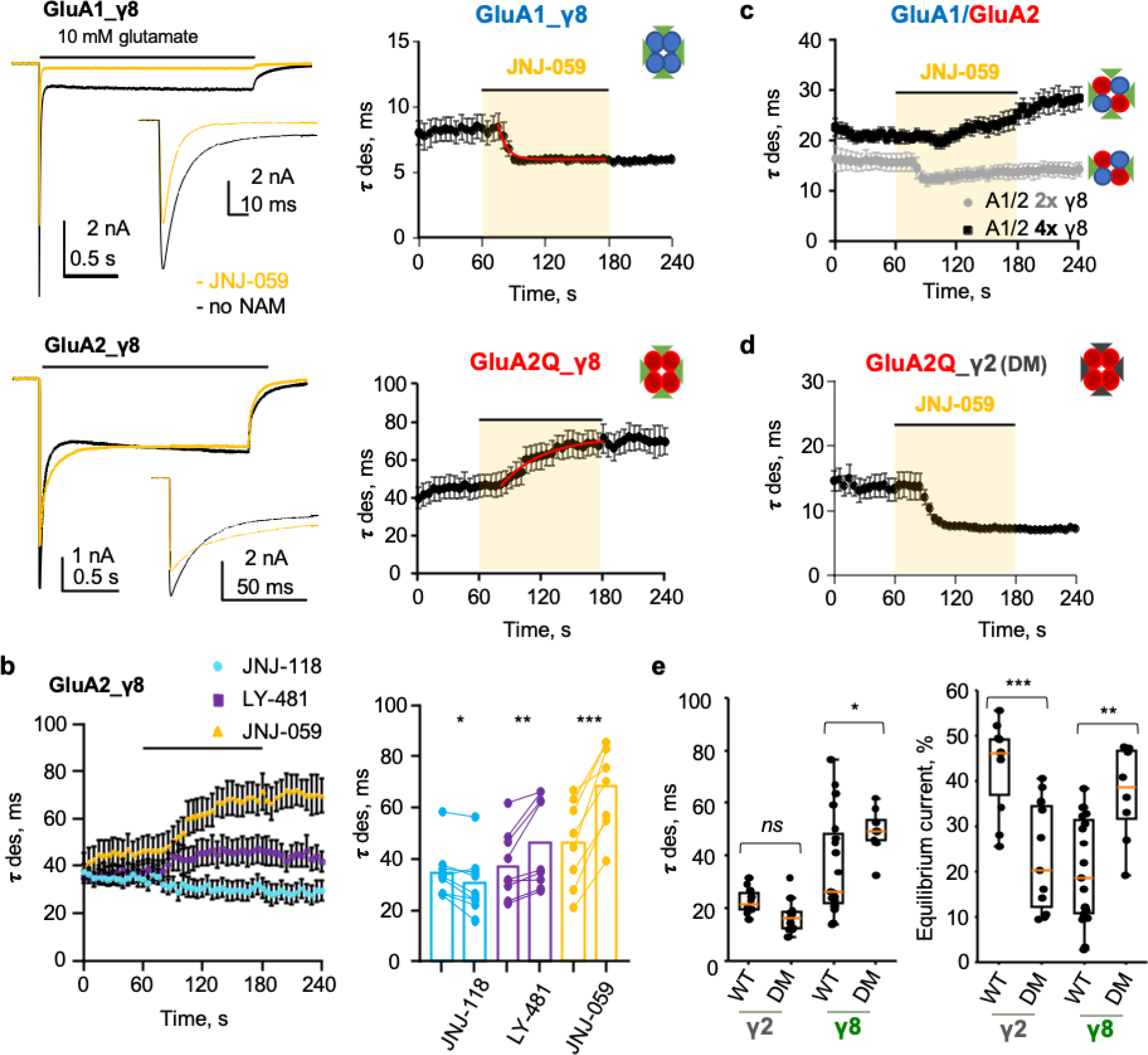
JNJ-059 PAM action on GluA2-containing AMPARs associated with four γ8 subunits. **a**, Left: Representative whole-cell responses to 10 mM glutamate (2 seconds, −60 mV) from HEK293T cells transfected with GluA1_γ8 (Top) or GluA2Q_γ8 (Bottom) tandem in control condition (black) and in presence of JNJ-059 10 μM (orange). Insets show initial phase of currents at faster time scale. Right: Scatter plot of average desensitization τ over time indicating the time-course of JNJ-059 effect on GluA1_γ8 (n=7; Top) or GluA2Q_γ8 (n=8; Bottom). Black circles and whiskers indicate average values and SD of the mean. Red line is a fit with single exponential. **b**, Left: Scatter plot of average desensitization τ in time indicating the time-course of modulator effect on GluA2Q_γ8. Black circles and whiskers indicate average values and SD of the mean. Number of cells are n=9 for JNJ-118, n=9 for LY--481 and n=8 for JNJ-059. Right: Bar and paired plots showing the effect of all three modulators on the desensitization τ for GluA2Q_γ8 (control 35±3 ms, JNJ-118 31±4 ms, n=9; control 38±4 ms, LY-481 46±6 ms, n=9; control 46±6, JNJ-059 69±6, n=8). Each data point is shown by black circles and bars represents mean values. Asterisks indicate summary of two-tailed paired t-test values **c**, Scatter plot of average desensitization τ in time indicating the time-course of JNJ-059 effect on GluA1/GluA2R_γ8 (n=4; grey circles) or GluA1/GluA2R_γ8/γ8 (n=10; black squares). **d**, Scatter plot of average desensitization τ in time indicating the time-course of JNJ-059 effect on GluA2Q_γ2DM (n=5). Black circles and whiskers indicate average values and SD of the mean. **e**, Box plots showing macroscopic desensitization τ (Left) and equilibrium current (Right) for GluA1i_γ2 or γ8 (wild type or double mutant). Each point for the desensitization τ plots was obtained by fitting the decaying phase of whole-cell currents with one or two exponentials (γ2: 23±2 ms, n=10; γ2DM: 17±2 ms, n=11; γ8: 37±4 ms, n=19; γ2: 49±3 ms, n=8). Equilibrium currents are 43±3% for γ2, 23±4% for γ2DM, 20±3% for γ8 and 37±4% for γ8DM. Boxplots are as described in Fig 4C. Asterisks indicate summary of two-tailed not-paired t-test values (see Fig. 4a for details).

The slower kinetics of PAM development (∼ 5-fold slower than NAM action; **Fig. 6a**, right panels) suggest that a different mechanism underlies its expression. In support of this, mutating the tyrosine on top of the M1 site in GluA2 (Y523A, the equivalent of GluA1 Tyr519) did not block PAM action (**Fig. S9c**), despite its strong prevention of NAM action in GluA1. The same held for GluA2 Met527A. While the equivalent mutation in GluA1 (M523A) robustly abolished NAM activity (**Fig. 4b** and **c**), GluA2 M527A did not blunt the GluA2 PAM phenotype (**Fig. S9c**), suggesting that other receptor elements are recruited to elicit positive modulation by JNJ-059.

Despite the presence of GluA2, PAM activity was not apparent in the GluA1/2_γ8 heteromer associated with two γ8 subunits. This receptor responded to JNJ-059 with a NAM phenotype, comparable to GluA1_γ8 (**Fig. 6c**). Interestingly, PAM action was observed when co-expressing additional γ8 with GluA1/2_γ8 (**Fig. 6c**). Therefore, contrary to NAM activity, population of all four TARP binding sites on the receptor is required for positive modulation of receptor kinetics. To determine if γ8 was an essential requirement, or whether any TARP could mediate positive modulation by JNJ-059, we created a NAM binding site on TARP-γ2 (I153V, A184G), by mutating the non-conserved residues to those of γ8 (**Fig. S4d**) ^31,32^. Surprisingly, GluA2 assembled with the γ2 double mutant (GluA2_γ2 DM) triggered a strong NAM response with JNJ-059 (**Fig. 6d**). Therefore, JNJ-059 PAM activity requires the presence of GluA2 and four TARP γ8 auxiliary subunits.

We note that the γ2 mutant, mimicking γ8 at the NAM binding site (I153V, A184G), was less potent in modulating the receptor (**Fig. 6e**). On the contrary, mutating γ8 at these two positions, to match the other Type-1 TARPs (V176I/G209A), converted γ8 into a more potent TARP (**Fig. 6e**). This result further demonstrates the fine balance in modulatory activity, which can be achieved by specific binding modes at the TARP-AMPAR interface around the NAM binding site. This strategic region has been exploited in different ways by other auxiliary subunits to fine-tune the AMPAR response across the brain.

## Discussion

Our study provides both an in-depth characterization and a mechanistic understanding of the modulatory action of TARP γ8 drugs. This novel class of compounds selectively targets TARP γ8-containing AMPARs and show promise in the treatment of epilepsy ^31,32^, pain and malignant gliomas ^33,34^. We demonstrate that their action is intricate and multi-modal, and is not simply due to an all-or-none occlusion of gate opening, as is the case for drugs targeting the AMPAR core (between pre-M1 and M3) ^43,44^. This is in line with their modulatory (rather than inhibitory) activity, and highlights the unique niche for targeting the receptor periphery for more subtle and specific action. The diversity of AMPAR auxiliary subunits provides ample opportunity for modulator development in future screening campaigns ^9^.

NAM binding to the AMPAR/TARP interface triggers both local and global effects. The NAM pocket borders the pre-M1 helix and M4 gating linkers, both poised to effectively transmit any local perturbation directly to the gate. This strategic location has been exploited by other auxiliary subunits. In CNIHs (CNIH1-3), three highly conserved phenylalanine side chains slot into a region overlapping the NAM pocket, and are central to positive modulation by CNIHs ^22,25^. Interestingly, the negative modulatory GSG1L subunit harbors a bulky tryprophan side chain at this position, at a site analogous to γ8 Gly209 (the γ8-selective NAM anchor point). The functional impact of this residue is currently unknown. Although the precise allosteric routes leading from these elements to the M3 gate are not yet firmly delineated multiple contacts of M3 with pre-M1 and the M4 linkers exist ^45,49^. For example, H-bonding of Tyr519 (in M1) to Asp515 (in pre-M1) in response to JNJ-059 could signal through Pro516, which is in van der Waals distance of Phe619 in the M3 gate (and is adjacent to Asp515 in pre-M1). This outlines one potential signaling route leading from the NAM pocket to the gate. The case for annular lipids as critical components of both drug and TARP action is building - they are consistently observed lining the side of the NAM pocket in AMPAR cryo-EM structrues ^10^, and mutation of lipid-interacting residues consistently affects NAM activity in this study. Lipid rearrangement in response to NAMs may not only shape the dynamics of pre-M1 but also bridge the NAM pocket with the helices deep in the conduction path (**Fig. S5g**).

Extracellular loops have been intensively studied in attempts to understand TARP modulation ^50,51^, yet our results now highlight the, currently understudied, role of the TARP transmembrane sector ^52^. Mutations thoughout M1 and M4 result in either positive or negative TARP action, implying that individual γ8 contact points are sensitive to modulation, and can be communicated differently to various points along the conduction path. The upper portion of M1 (encompassing the NAM binding site) faces pre-M1 and the M3 gate and is more loosely connected to γ8 ^38^, while the more tightly connected lower half of M1 is level with the selectivity filter (**Fig. 5d**). Interestingly, alanine mutations in the more tightly coupled lower segment generally convey positive TARP action while those at the upper M1 end are either neutral or negative. Closer inspection also reveals that alanine mutants have differential impacts on various components of the TARP modulatory spectrum; for example, despite slowing kinetics, F537A does not markedly affect the equilibrium response, while the opposite appears to hold for L785A (**Fig. 5** and **Fig. S8a**). Moreover, R541A severely slows desensitization entry but completely blocks resensitization, thus positively modulating one functional component but blunting another. It is upon this finely balanced receptor-TARP interaction landscape that NAMs act.

We propose that existing γ8 NAMs could be exploited further through targeted drug design in future efforts. Exposure of the NAM pocket to lipid ^53^, together with unoccupied cavities, provide an opportunity for continued development of novel modulators. For example, in the LY-481 structure, the pocket extends from the oxindole carbonyl toward Ser128 at the kink of the γ8 M2 helix. This region is highly electropositive and accessible to water, with putative water molecules coordinated by the Ser128 side chain (**Fig. S4c and 5g**). In fact, the less bulky tryptophan metabolites isatin and 5-OH-indole penetrate deeper into this space, and thus might lead the way as starting points for drug design (**Fig. S7b**).

Novel modulators will not only include NAM action.The unexpected JNJ-059 PAM response with GluA2 appears to engage a different allosteric route. Expression of the PAM phenotype is approximately 5-fold slower than the NAM effect, and is insensitive to mutation at Tyr523 and Met527, while the equivalent positions in GluA1 blunt JNJ-059 NAM action. Together with a strict requirement for four γ8 subunits, and the inability of γ2 to mediate the effect, we hypothesize that the extracellular, sequence-diverse TARP loops are involved. These project towards the LBDs, which, contrary to the TMD sector, exhibit sequence differences between GluA1 and GluA2. Interaction between these loops and the AMPAR LBDs is gating state-dependent ^38^, and is known to regulate desensitization kinetics ^45,50,51^. Moreover, TARPs at either the A’C’ or B’D’ sites will have a different reach for the LBDs and the gating linkers ^19^. The requirement for four TARPs suggests that A’C’ interactions are essential for positive modulation. One fascinating aspect is why specifically JNJ-059 triggers this response. Further derivatives of this compound may lead to improved PAMs, and thus potentially to drugs that selectively boost cognition in the hippocampus ^54^.

## Acknowledgments

We thank James Krieger for generating the ‘proDy’ interaction maps in Figure 5B and S7C, and Jan-Niklas Dohrke for critically reading the manuscript. We thank members of the Greger lab for insightful comments during this study. We acknowledge Trevor Rutherford for confirming ligand integrity by NMR. We are also grateful to LMB scientific computing and the EM facility for their support. This work was supported by the Medical Research Council, as part of United Kingdom Research and Innovation (also known as UK Research and Innovation) MRU105174197] and the Wellcome Trust [223194/Z/21/Z] to I.H.G, and NIH grant (R56/R01MH123474) to T.N. For the purpose of open access, the MRC Laboratory of Molecular Biology has applied a CC BY public copyright licence to any Author Accepted Manuscript version arising.

## Author contribution

I.H.G supervised the study, and wrote the paper with input from D.Z. R.L., S.A.S, J.F.W. and T.N. D.Z. and B.K.K. performed protein purification and cryo-EM data collection, D.Z. and B.K.K. performed data processing with input from T.N. R.L. and O.C. designed and performed electrophysiological experiments, J.F.W. contributed key observations at early stages of this study, S.A.S performed and analysed MD simulations.

## Declaration of interests

The authors declare no competing interests.

## Materials and Methods

### cDNA constructs

All cDNA constructs were produced using IVA cloning (Garcia-Nafria et al., 2016). Constructs used for structural studies in this paper are the same as previously reported, and are all cloned into pRK5 vectors. To express GluA1/A2_γ8 recombinantly, GluA1 (rat cDNA sequence, flip isoform) was fused with a FLAG tag at the N-terminus, and GluA2 (rat cDNA sequence, V1 to S839, flip isoform, R/G edited, Q/R edited) was cloned in a tandem configuration with a GGSGSG linker to TARP γ8 (rat cDNA sequence, E2-K419), a human rhinovirus 3C (HRV 3C) protease cleavage site and an eGFP.

### Expression and purification of GluA1/2_γ8

To achieve heteromeric AMPAR expression, FLAG-tagged GluA1 and GluA2- TARPγ8-eGFP tandem plasmids were co-transfected into HEK-Expi293™ cells at a ratio of 1:1. To prevent AMPA-mediated excitotoxicity, AMPAR antagonists ZK200775 (2 nM, Tocris, Cat# 2345) and kynurenic acid (0.1 mM, Sigma, Cat# K335-5G) were added to the culture medium. 36-44 hours post-transfection, cells were harvested and lysed for 3 hours in lysis buffer containing: 25 mM Tris pH 8, 150 mM NaCl, 0.6 % digitonin (w/v) (Sigma, Cat# 300410-5G), 5 μM NBQX, 1 mM PMSF, 1× Protease Inhibitor (Roche, Cat# 05056489001). Insoluble material was then removed by ultracentrifugation (41,000 rpm, 1 hour, rotor 45-50 Ti) and the clarified lysate incubated with anti-GFP beads for 3 hours. After washing with glyco-diosgenin (GDN) (Anatrace, Cat# GDN101) buffer (25 mM Tris pH 8, 150 mM NaCl, 0.02% GDN) the protein was eluted from the beads by digestion with 1 mg/ml 3C protease at 4 °C overnight. Eluted fractions were incubated with ANTI-FLAG M2 affinity gel (Sigma, Cat# A2220) for 1.5 hours and washed 3 times with GDN buffer. Finally, the complex was eluted using 0.15 mg/ml 3×FLAG peptide (Millipore Cat# F4799) in GDN buffer. Eluted fractions were pooled and concentrated to ∼2.5 mg/ml for cryo-EM grid preparation.

### Cryo-EM grid preparation and data collection

Cryo-EM grids were prepared using a FEI Vitrobot Mark IV. For the resting state GluA1/A2_γ8 heteromeric complex, protein was incubated with 300 μM ZK200775 and 40 μM ligands (JNJ-61432059, MedChemExpress, Cat# HY-111751, JNJ- 55511118, Tocris, Cat#6278, LY-3130481, MedChemExpress, Cat# HY-108707) for at least 30 min on ice before freezing. For the active state A1/A2_γ8/C2 heteromeric complex, protein was first incubated with 300 μM cyclothiazide (CTZ, Tocris, Cat# 0713) and 40 μM JNJ-059 for at least 30 min on ice and then quickly mixed with 1 M L-glutamate stock solution to a final concentration of 100 mM prior to loading onto the grids. Quantifoil Au 1.2/1.3 grids (300 mesh) were glow-discharged for 30 s at 0.35 mA before use. 3 μl sample was applied to the grids, blotted for 4.5-5 s at 4 °C with 100 % humidity and plunge-frozen in liquid ethane. All cryo-EM data were collected on a FEI Titan Krios operated at 300 kV, equipped with a K3 detector (Gatan) and a GIF Quantum energy filter (slit width 20 eV). Movies at 1.5-2.5 μm underfocus were taken in counting mode with a pixel size of 1.07 Å/pixel. A combined total dose of 50 e/Å^2^ was applied with each exposure and 50 frames were recorded for each movie.

### Cryo-EM data processing and model building

Dose-fractionated image stacks were first motion-corrected using MotionCor2 ^57^. Corrected sums were used for CTF estimation by GCTF ^58^. All further data processing was performed with RELION 3.1 ^59^. Automatic particle picking was performed using a Gaussian blob and particles were binned to 4.28 Å/pixel and extracted in a box of 80 pixels. 2 to 3 rounds of 2D classification were carried out to remove particles not showing AMPAR-like features. For the following 3D classification, emd-4575 was used as initial model to further eliminate low-quality particles. Following data clean-up, particles were re-centered, scaled up to 2.14 Å/pixel and re-extracted in a box of 160 pixels. Another 3D classification was applied on the re-extracted data set. After this round of classification, selected particles were scaled to the original 1.07 Å/pixel size, and refined with C1-symmetry followed by post-processing. CTF refinement and Bayesian polishing were then performed, followed by another refinement. To further improve map resolution, we applied masked refinement on LBD-TMD region and TMD region alone with C2-symmetry introduced. Local resolution was estimated by RELION3.1.

Model building and refinement were performed using Coot ^60^, REFMAC5 ^61^ and PHENIX ^62^ real-space refinement. The GluA1/A2-γ8 complex (PDB 6QKC) was used as starting points, and rigid-body fitted into the maps using UCSF chimera (http://www.rbvi.ucsf.edu/chimera) then automatically refined by REFMAC5. Afterwards, manual refinement was performed through Coot, followed by PHENIX real-space refinement to further refine the geometry. Ligand models were first built manually in Coot by using ligand builder and then fitted into maps. Corresponding restraint files were generated by using PHENIX elbow. After merging ligand and refined protein coordinates, final refinements were carried out by using PHENIX real-space refinement and model validation was performed with MolProbity ^63^. LY-481 was built as (-) enantiomer based on a previous study (Gardinier *et al*., 2016). Graphics were prepared using UCSF Chimera, ChimeraX or PyMOL (http://www.pymol.org). Pore radius was calculated using a plugin version of HOLE ^64^ in Coot.

### All-atom MD simulations

Four ligand -bound, and two apo structures of the LBD-TMD-TARP complex described in this and previous studies from the lab ^38,65^, were used to set up simulations. Missing residues and loops were added using MODELLER ^66^, with restraints applied to the β1-loop to extend its conformation. 100 models were generated, and three models with the highest consensus DOPE score ^67^ and SOAP- LOOP score ^68^ were used for simulation. CHARMM-GUI ^69^ was used for system setup. TARP residues N53 and N56 were glycosylated, and TARP E216 was protonated as it is surrounded by hydrophobic residues. CGENFF parameters ^70^ were used for all ligands. The protein complexes were embedded in POPC lipid bilayers. TIP3P water was used to solvate the system and 150 mM NaCl was added.

Three simulations were performed for each of the six systems. After minimization for 10000 steps, two equilibration steps in the NVT ensemble of 125 ps each, an equilibration step in the NPT ensemble of 125 ps with a 1 fs time step, followed by three equilibration steps in the NPT ensemble, each 500 ps with a 2 fs time step, were performed, where harmonic restraints on the protein, and planar/dihedral constraints on the lipids were consecutively decreased. Following removal of all restraints, the NPT ensemble was used for production runs. All simulations completed 500 ns, except one of the repeats for JNJ-118 that we had to end at 250 ns due to compute resource constraints; we thus obtained a cumulative of 1.25 microseconds of sampling for JNJ-118, and 1.5 microsecond sampling for all other systems. These simulations were performed using NAMD 3.0 alpha ^71^, where the simulation temperature was controlled at 303.15 K by Langevin dynamics, with a damping coefficient of 1 ps^−1^, and the pressure of the system was kept at 101.325 kPa (1.01325 bar or 1 atmosphere) using the Nosé-Hoover Langevin method ^72^ with a piston period of 200 fs and piston oscillation decay time of 100 fs. The CHARMM36m force-field ^73^ and a 2.0 fs time step for production runs was used for all systems. Analysis of the simulation data and preparation of graphics was done using VMD 1.9.3 ^74^. All analyses were performed with a sampling of 100 ps/frame. Contact maps were generated by calculating the percentage of simulation time unique inter-residue contacts existed, determined as residues having at least one non-H atom each within a distance of 4 Å of each other, using custom scripts with VMD 1.9.3. The first 100 ns of each simulation was excluded for contacts calculations to allow an extended equilibration time, and resulting percentages were averaged over identical subunits and all three repeat runs. To generate difference maps, the difference in contact percentages between apo and ligand- bound states in the same state, i.e., open vs open and resting vs. resting were taken, excluding any contact percentage < 10% as well as difference < 10%, to reduce noise.

A single set each of 350 ns long simulations was also performed for only the transmembrane domains of the A1/A2/g8 complex with JNJ059, isatin, and 5- hydroxyindole at the JNJ059 site (the latter two were modeled in, based on the JNJ059 oxindole position). The simulation protocol followed was the same as described above for ligand bound complexes.

### Patch-clamp recording

Glutamate-evoked currents were recorded from transfected HEK293T cells in the whole-cell or outside-out patch-clamp configuration at room temperature. The cells were bathed in the extracellular solution consisting of (in mM): NaCl (145), KCl (3), CaCl2 (2), MgCl2 (1), glucose (10), and HEPES (10), adjusted to pH 7.4 using NaOH. Recording pipettes were pulled from thick-walled borosilicate glass capillary tubes (1.5mm OD, 0.86mm ID; Science Products GmbH) on a Flaming/Brown puller P-1000 (Sutter Instruments), fire-polished to give a final resistance of 3-6 MΩ and filled with the intracellular solution containing (in mM): CsCl (130), EGTA (10), ATP- sodium salt (2), HEPES (10), and spermine (0.1), adjusted to pH 7.3 with CsOH for whole-cell recordings (junction potential +5.3 mV). For outside-out recordings the pipette solution contained 120 mM CsF and 10 mM CsCl (junction potential + 9.1 mV) instead of 130 mM CsCl. Recordings were performed by applying L-glutamate (10 mM) onto lifted cells or outside-out patches held at -60 mV transmembrane potential (not corrected for the junction potential). Access resistance in whole-cell recordings was monitored but not compensated. Cells with access resistance unstable or higher than 20 MΩ were excluded from the analysis. Agonist was dissolved in extracellular solution and applied using a double-barrel pipette (#BT- 150-10; Science Products GmbH) mounted on a piezo actuator (Physik Instrumente). Data were acquired and filtered (at 5 kHz 4-pole Bessel filter) with MultiClamp 700B amplifier (Molecular Devices), digitized with Digidata 1440A (Molecular Devices) at a sampling rate of 20 kHz and saved into PC.

In the whole-cell recordings glutamate was applied to the cells in 2 s pulses every 5 s. Control currents were recorded for at least 60 s then tested compounds were applied during and between glutamate pulses for 120 s and then washed off. Coverslips with the cells were removed after each application of compound to avoid recording from cells pre-exposed to compound. The time-course of macroscopic desensitization was measured by fitting the decay phase of the currents from 95% to the steady-state current with one or two exponentials. Equilibrium current was measured as steady-state current percentage of peak current. The resensitization was defined as an excess steady-state current following desensitization through to the end of 2 s glutamate pulse and expressed as the percentage of peak current. Changes in peak current were quantified by measuring and averaging (5 sweeps) peak amplitude just before the application of the ligand and at the full extent of the modulatory effect- at 1 min or 2 min after ligand application for negative or positive modulation, respectively. Modulatory compounds were dissolved in DMSO to 50 mM as stock solution and used at final concentration of 10 μM.

### Quantification and statistical analysis

Summary data are presented as mean ± SD of the mean. Statistical tests were performed using GraphPad Prism 9.4.0. *p* values were calculated from one- or two-sample (paired or unpaired) two-tailed t-tests. Ordinary one-way ANOVA test with Dunnett correction was used for multiple comparisons. *p* values in the figures are indicated as * for *p*<=0.05, ** for *p*<=0.01, *** for *p*<= 0.001 and ‘ns’ for *p*>0.05.

## Notes

### Competing Interest Statement

The authors have declared no competing interest.

