## Supplemental Material for "Modulatory mechanisms of TARP γ8-selective AMPA receptor therapeutics"

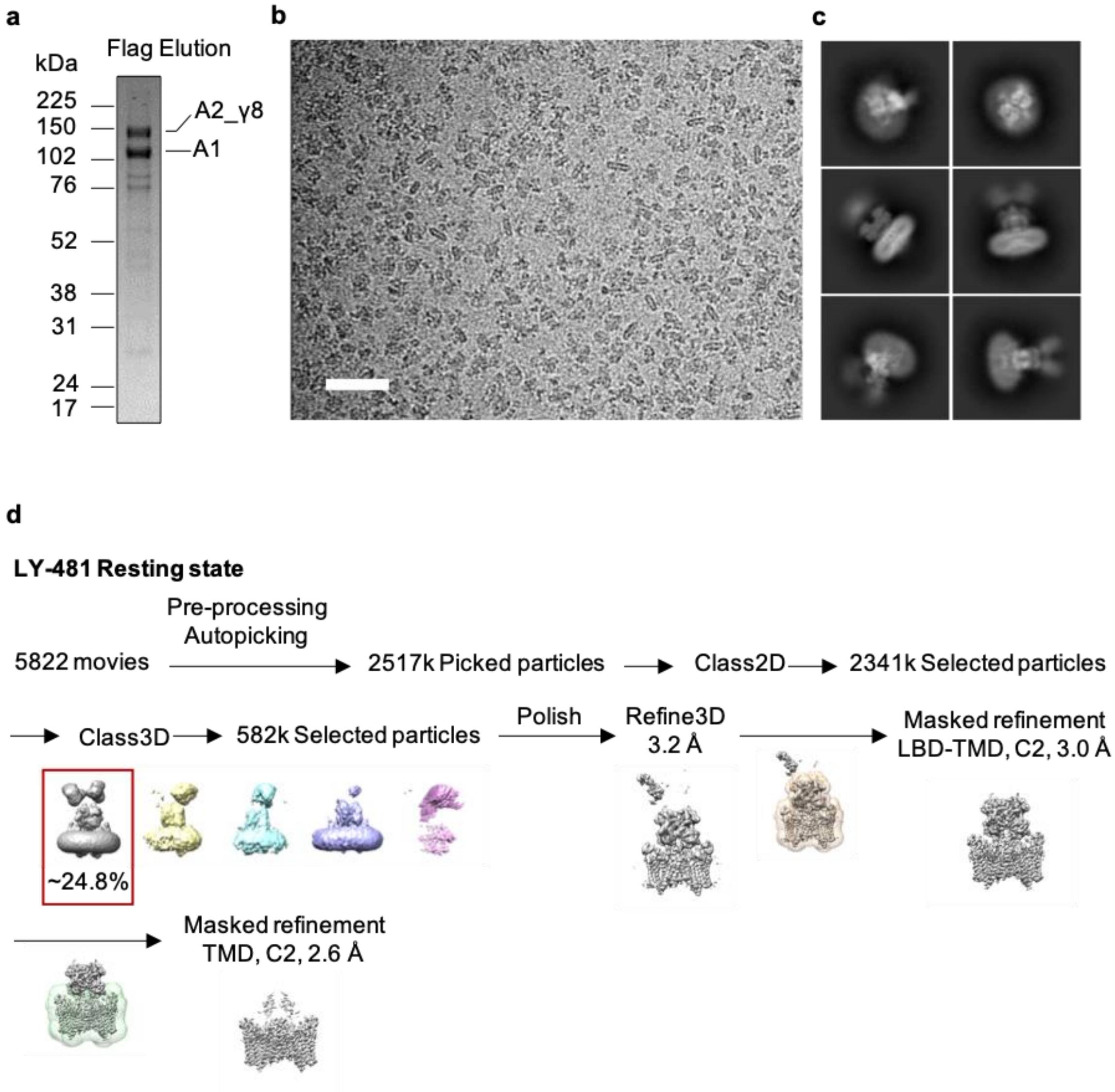


**Fig. S1. Purification of recombinant AMPAR complex and representative Cryo-EM data processing workflow of GluA1/A2_γ8 in complex with γ8 NAMs.**

**a**, Representative 4-12% Bis-Tris SDS-PAGE gel stained with Coomassie blue, indicating elution of the GluA1/2_γ8 complex from FLAG beads. **b**, Representative, motion-corrected micrograph of resting-state GluA1/2_γ8 in complex with LY-481 (scale bar, 50 nm). **c**, Representative 2D class averages of the resting state GluA1/2_γ8 in complex with LY-481. **d**, Cryo-EM data processing workflow of the resting state GluA1/2_γ8 in complex with LY-481. Raw movies were first processed, then more than 2 million raw particles were picked from motion-corrected micrographs. Then 2D and 3D classification was performed to remove bad particles, and finally 582k particles were selected and polished for refinement. Next, focused refinement on LBD-TMD gating core was performed with C2 symmetry. To further improve the resolution at the ligand-binding pocket, the TMD sector alone was refined with C2 symmetry applied. This workflow was also implemented for the other structures (JNJ-118, JNJ-059).

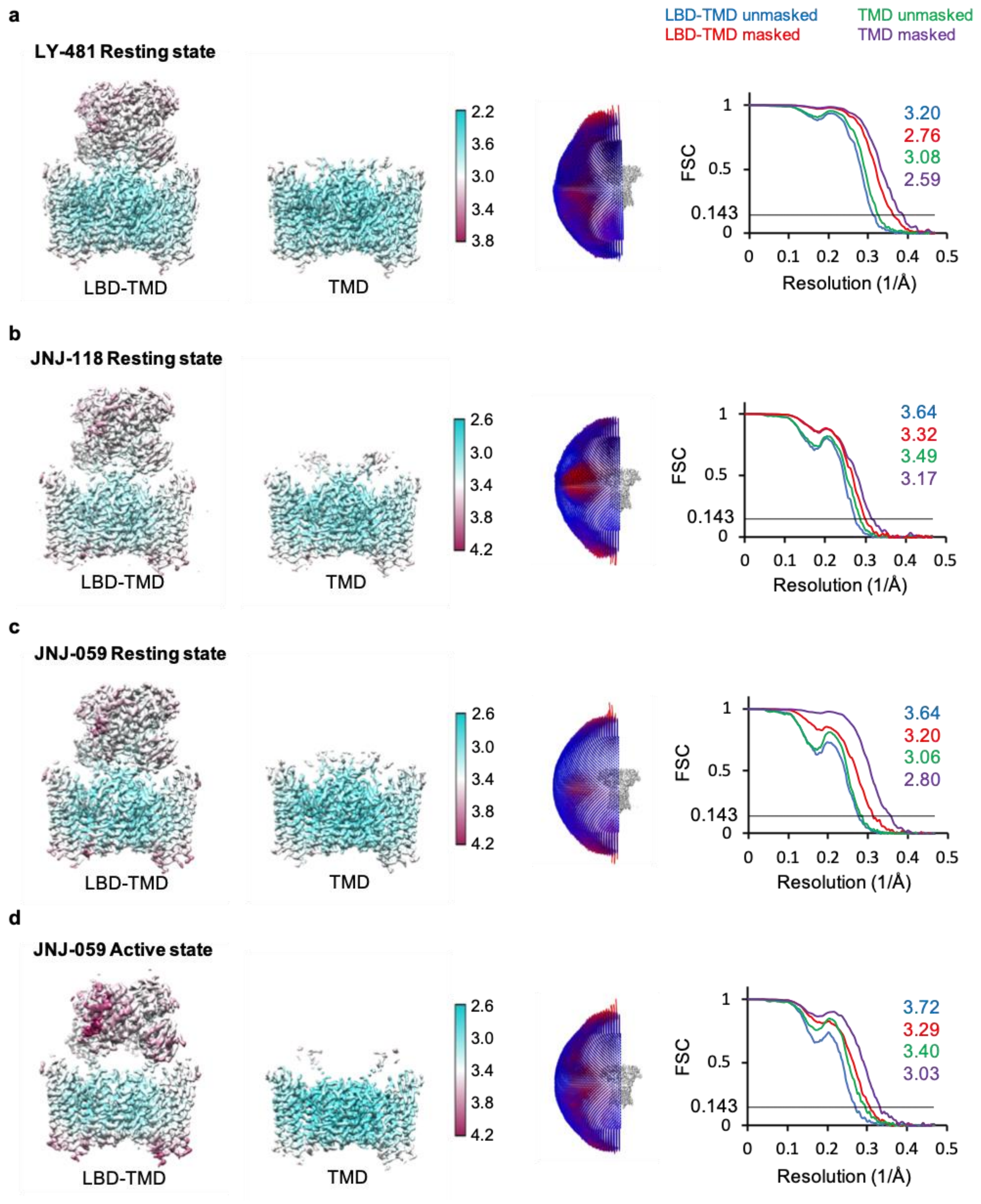


**Fig. S2. Cryo-EM analysis of GluA1/A2_γ8 in complex with γ8 NAMs.**

**a**, **Left:** Local resolution maps at LBD-TMD and TMD of resting state GluA1/A2_γ8 in complex with LY481. 3D maps are coloured based on local resolution estimate. **Middle:** Euler angle distribution of particles used for the cryo-EM reconstruction. **Right:** Masked (red, LBD-TMD and purple, TMD) or unmasked (blue, LBD-TMD and green, TMD) Fourier shell correlation (FSC) of corresponding maps where FSC=0.143

(black line). **b**, Local resolution maps, particle Euler angle distribution and FSC curves of resting state GluA1/A2_γ8 in complex with JNJ-118. Figures are coloured as in A. **c**, Local resolution maps, particle Euler angle distribution and FSC curves of resting state GluA1/A2_γ8 in complex with JNJ-059. Figures are coloured as in A. **d**, Local resolution maps, particle Euler angle distribution and FSC curves of open state GluA1/A2_γ8 in complex with JNJ-059. Figures are coloured as in panel A.

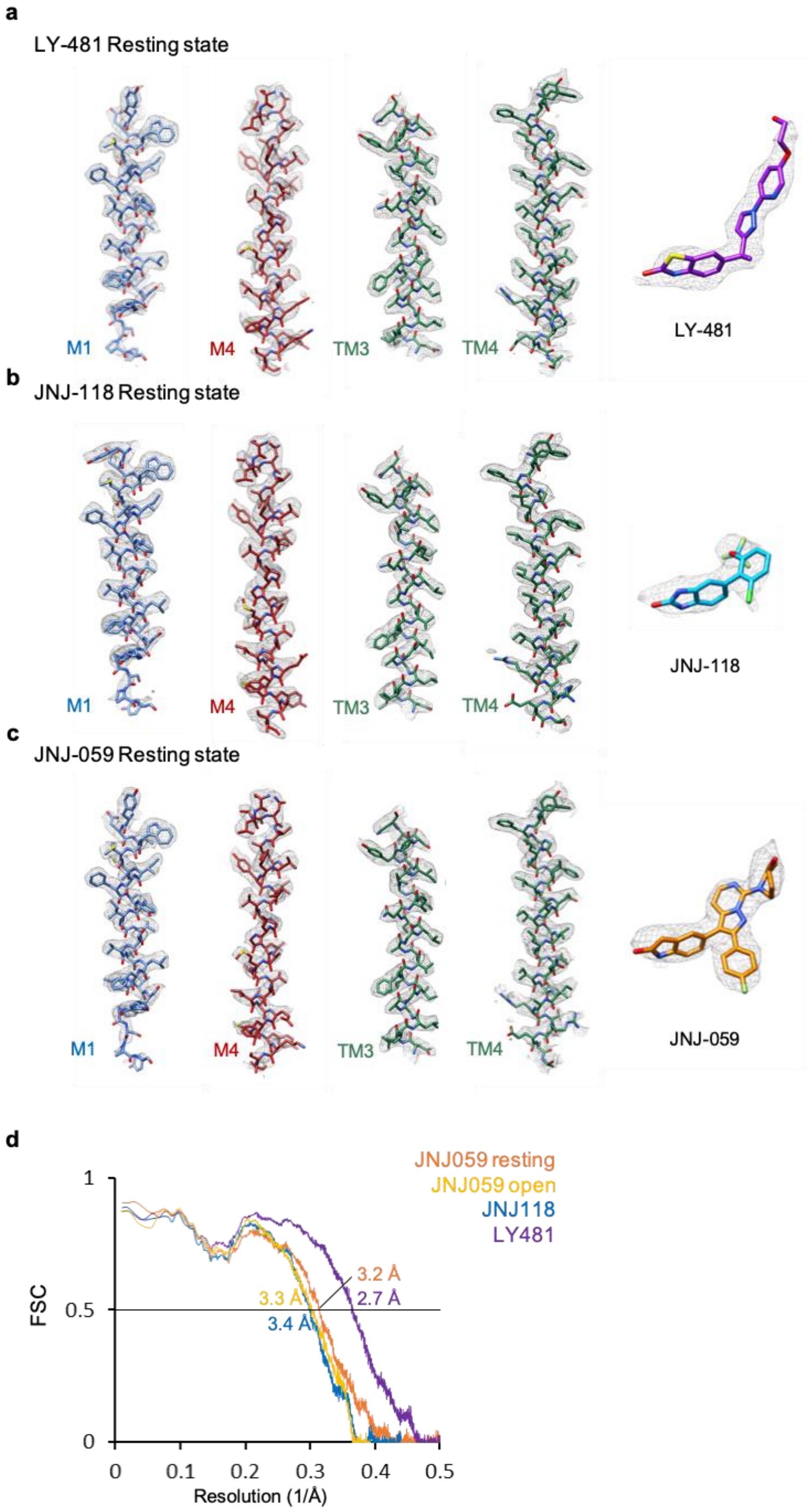
a
LY-481 Resting state
M1
M4
TM3
TM4
LY-481
b
JNJ-118 Resting state
M1
M4
TM3
TM4
JNJ-118
c
JNJ-059 Resting state
M1
M4
TM3
TM4
JNJ-059
d
JNJ059 resting
JNJ059 open
JNJ118
LY481
FSC
Resolution (1/Å)
3.2 Å
2.7 Å
3.3 Å
3.4 Å
1
0.5
0
0.1
0.2
0.3
0.4

**Fig. S3. Densities and their fit against models of GluA1/A2_γ8 transmembrane helices.**

**a**, Densities of LY-481 and surrounding transmembrane helices M1(GluA1), M4(GluA2), M3(γ8), M4(γ8) and their fit against the model. **b**, Densities of JNJ-118 and surrounding transmembrane helices M1(GluA1), M4(GluA2), M3(γ8), M4(γ8) and their fit against the model. **c**, Densities of JNJ-059 and surrounding transmembrane helices M1(GluA1), M4(GluA2), M3(γ8), M4(γ8) and their fit against the model. **d**, Model-to-map FSCs of ligand-bound GluA1/2_γ8 LBD-TMD models in resting and active states.

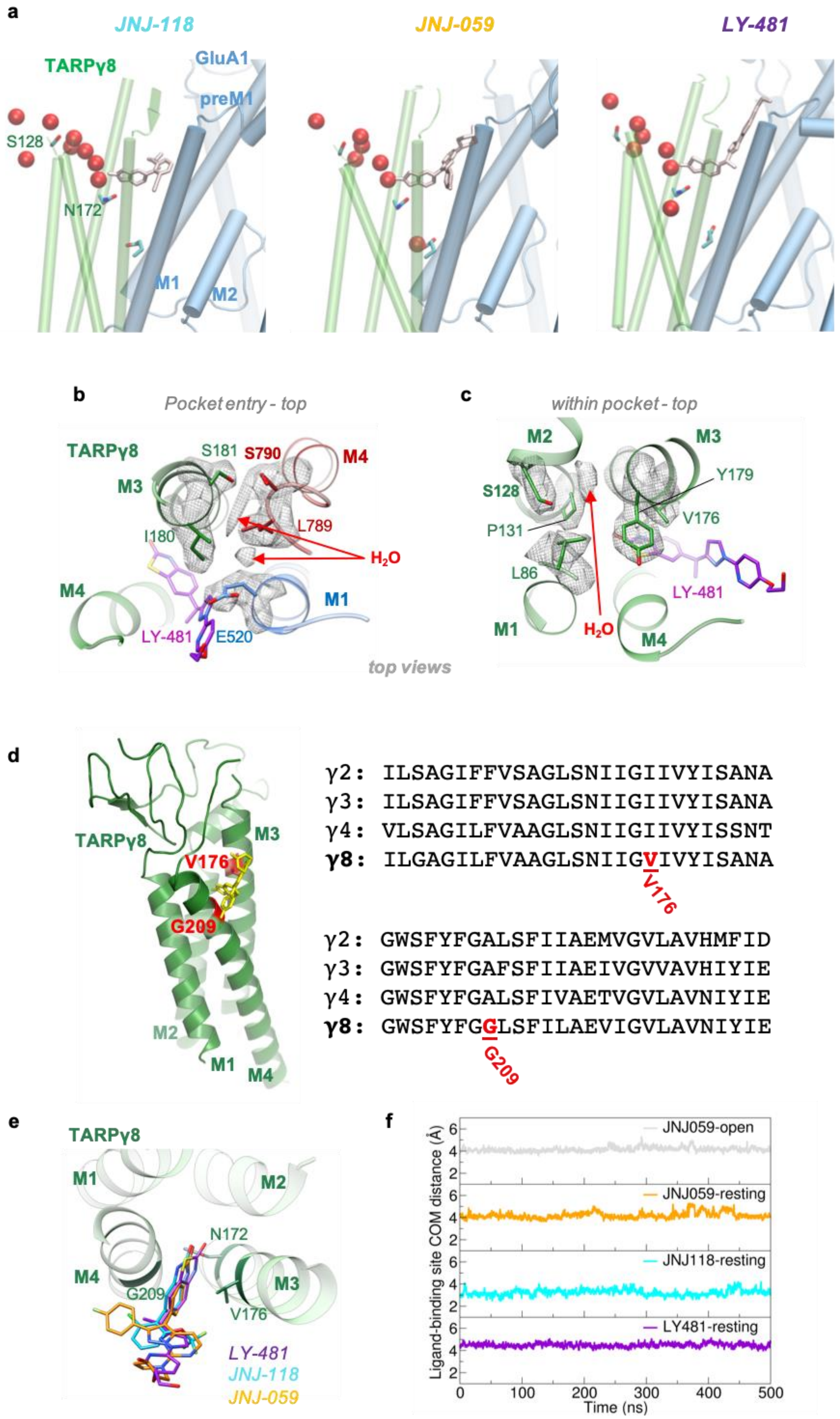
a
JNJ-118
JNJ-059
LY-481
TARPγ8
GluA1
preM1
S128
N172
M1
M2
b
Pocket entry - top
TARPγ8
S181
S790
M4
M3
L789
I180
$H_2O$
M4
M1
LY-481
E520
top views
c
within pocket - top
M2
M3
S128
Y179
P131
V176
L86
LY-481
M1
$H_2O$
M4
d
TARPγ8
M3
V176
G209
M2
M1
M4
γ2: ILSAGIFFVSAGLSNIIGIIVYISANA
γ3: ILSAGIFFVSAGLSNIIGIIVYISANA
γ4: VLSAGILFVAAGLSNIIGIIVYISSNT
γ8: ILGAGILFVAAGLSNIIGVIVYISANA
V176
γ2: GWSFYFGALSFIIAEMVGVLAVHMFID
γ3: GWSFYFGAFSFIIAEIVGVVAVHIYIE
γ4: GWSFYFGALSFIVAETVGVLAVNIYIE
γ8: GWSFYFGGLSFILAEVIGVLAVNIYIE
G209
e
TARPγ8
M1
M2
N172
M4
M3
G209
V176
LY-481
JNJ-118
JNJ-059
f
Ligand-binding site COM distance (Å)
JNJ059-open
JNJ059-resting
JNJ118-resting
LY481-resting
Time (ns)
0
100
200
300
400
500

**Fig. S4. Features of the NAM binding sites from cryo-EM and MD simulation analysis.**

**a**, Snapshots from MD simulations (left: JNJ-118, centre, JNJ-059, right: LY-481) showing water molecules from the extracellular side penetrating into TARP in the direction of S128 to N172 at the ligand binding site. **b**, Density of putative waters (indicated by the red arrow) and surrounding residues in the LY-481 cryo-EM map around GluA2 S790. Model is coloured as in Fig2. **c**, Density of putative waters (indicated by the red arrow) and surrounding residues in the LY-481 cryo-EM map around γ8 S128. Model is coloured as in Fig2. **d**, Sequence alignment of Type 1 TARPs, γ8-specific V176 and G209 are highlighted. **e**, Overlay of resting state LY-481, JNJ-118 and JNJ-059 models. Models are coloured as in Fig2. **f**, Ligand stability in the binding site, measured as the centre of mass (COM) distance of ligand heavy atoms from COM of C$\alpha$ of binding site residues, V176, G209, M523 and C524. Representative variations from one binding site in one simulation set for each system are shown. Low variation indicates the ligands remain bound in the site during simulations.

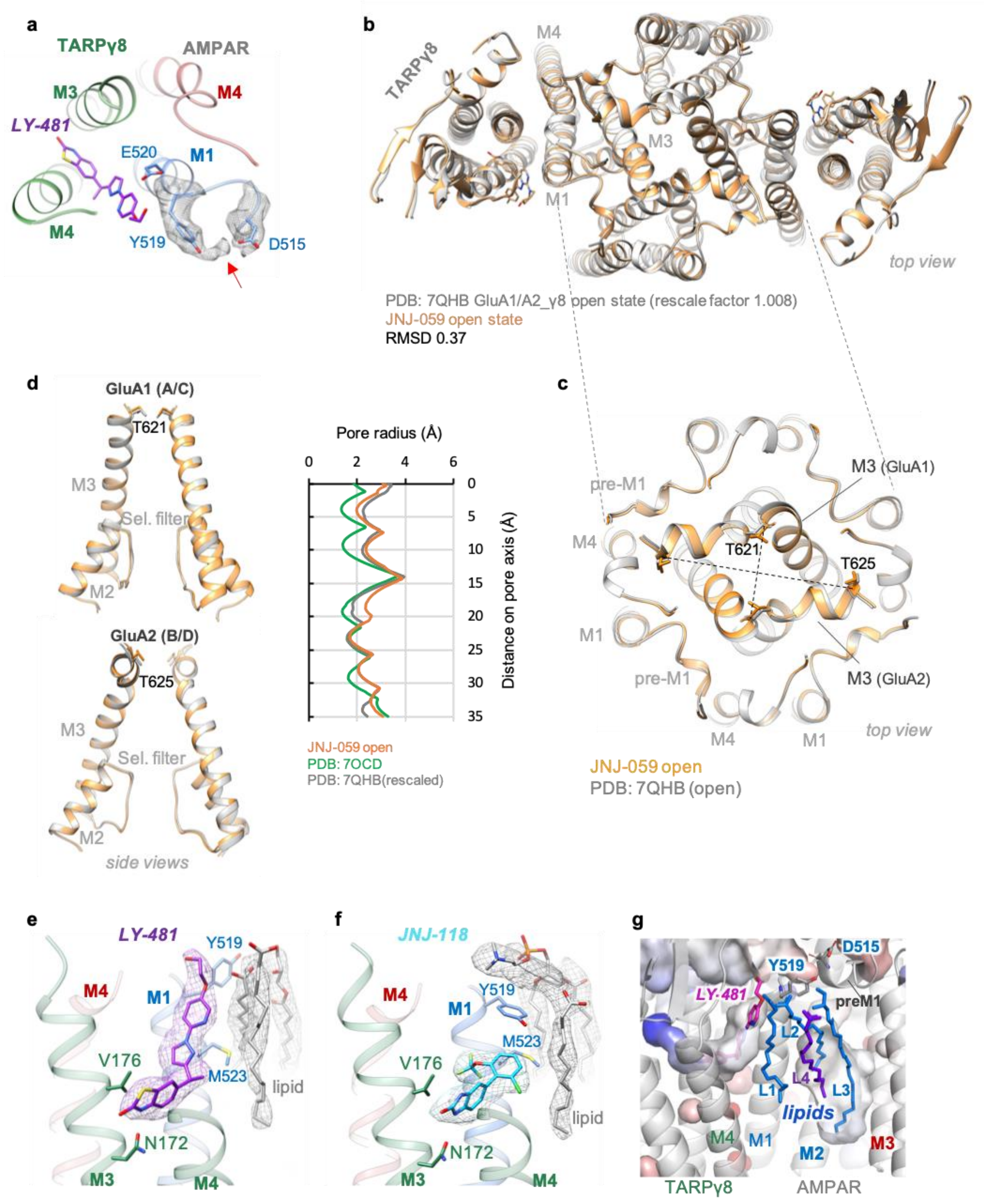


**Fig. S5. JNJ-059 open state and annular lipids.**

**a**, Density of putative water that bridges Y519 and D515 in the LY-481 cryo-EM map. Model is coloured as in Fig2. **b**, Superposition of open state GluA1/A2_γ8 in complex with JNJ-059 (orange) and the open state GluA1/A2_γ8 without ligand binding (PDB 7QHB; the model was rescaled by a factor of 1.008 to eliminate systematic error caused by pixel size calibration yielding better comparability). **c**, Zoomed-in view of the superposed models shows subtle changes at the receptor gate with JNJ-059 bound. **d**, Pore dimensions of resting state GluA1/A2_γ8 (green, PDB 7OCD) open state GluA1/A2_γ8 (grey, rescaled 7QHB as in panel B), and open state GluA1/A2_γ8 in complex with JNJ-059 (orange) depicted by space-filling representation. Side views of superposed M3 helices (top: GluA1, bottom: GluA2) from open state models are shown. **e**, Density of LY-481 and the lipid molecules surround ligand-binding pocket. Model is coloured as in Figure2. **f**, Density of JNJ-118 and lipid molecules surround the ligand binding pocket. Model is coloured as in Fig2. **g**, Cavities of the NAM (LY-481 serves as example) and annular lipids lining the conduction path helices (M2 and M3).

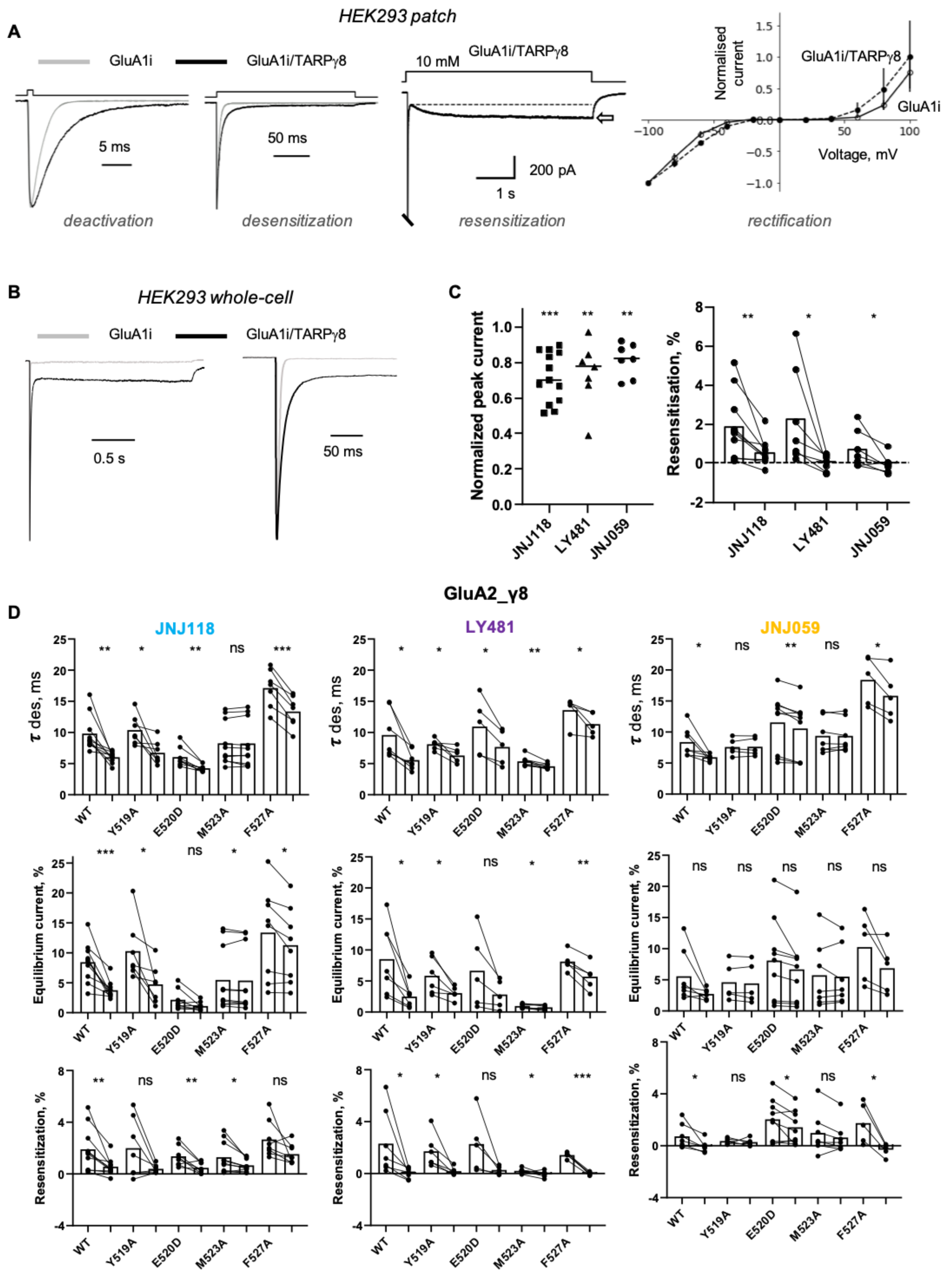


**Fig. S6. Electrophysiological characterisation of GluA1 wt and mutants in response to the three NAMs.**

**a**, Representative outside-out patch responses to 2 ms, 200 ms and 5 s pulses of 10 mM glutamate (transmembrane potential –60 mV) from HEK293T cells transfected with GluA1i alone (grey traces) or with TARP γ8 (black traces). Right: pooled normalized current-voltage relationships for GluA1i alone (n=6; open circles, black line) or with TARP γ8 (n=8; black circles, dashed line). **b**, **Left**: Representative whole-cell responses to 10 mM glutamate (2 seconds, –60 mV) from HEK293T cells transfected with GluA1i alone (grey trace) or GluA1i_γ8 tandem (black trace). **Right**: same traces shown at faster time scale. **c**, **Left**: Pooled scatter plot of GluA1i_γ8 current peak inhibition by three modulators. Each point is current peak in the presence of modulator normalized to the control peak. Horizontal lines indicate the mean values. **Right**: Paired plots showing effect of 10 µM JNJ-118, LY-481 or JNJ-059 on resensitization. **d**, Paired bar plots showing effect of 10 µM JNJ-118, LY-481 or JNJ-059 on desensitization $\tau$, equilibrium current and resensitization for wild type or mutated GluA1i_γ8. Each point is a measure of parameter in absence or presence of modulator. Bar height represents the mean value. Asterisks indicate summary of two-tailed paired t-test values: * $p <= 0.05$, ** $p <= 0.01$, *** $p <= 0.001$ and 'ns' for $p>0.05$. These data are represented in a different format in Fig 4c.

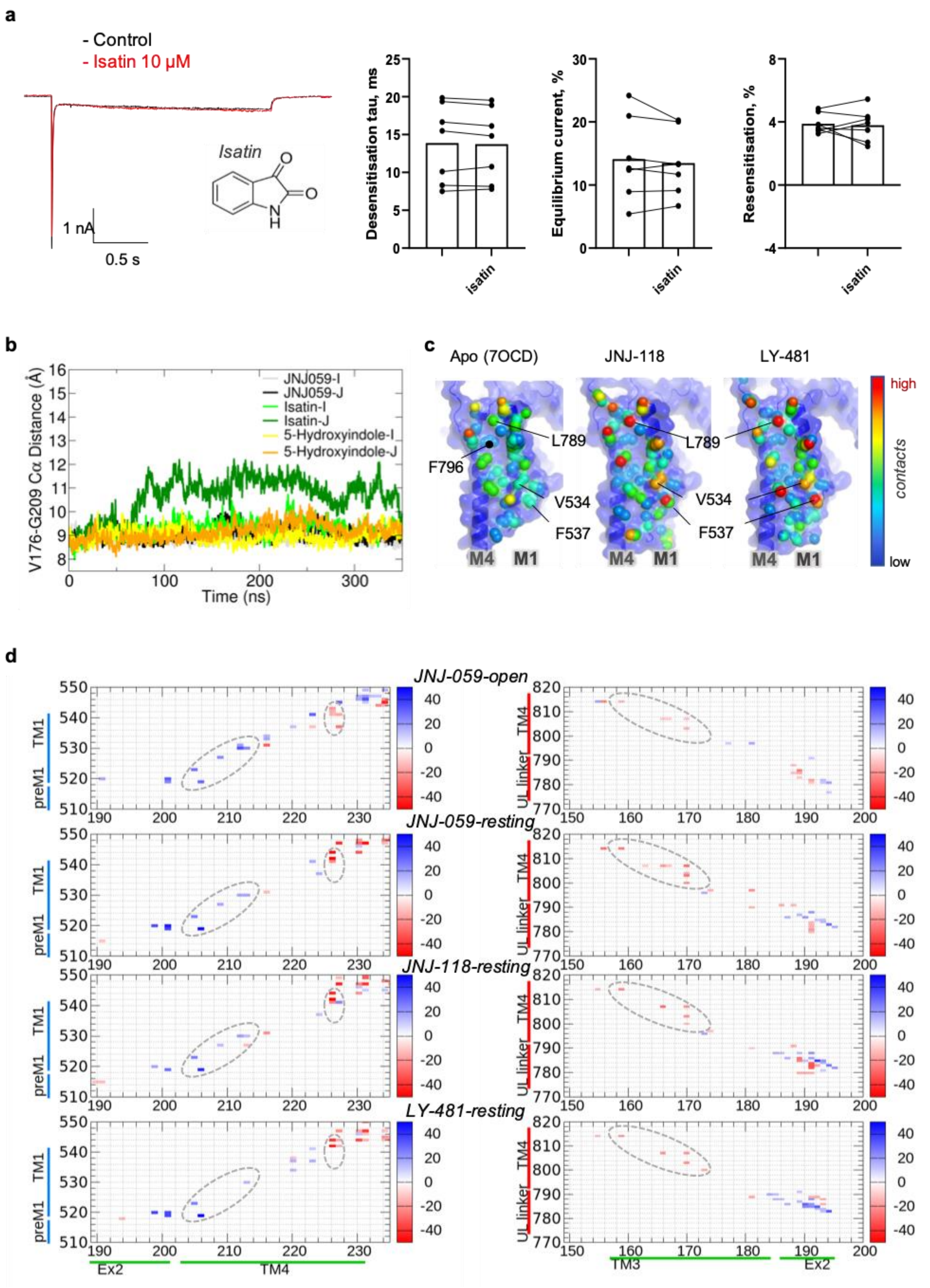


**Fig. S7. Role of natural oxindoles and contact map analyses.**

**a**, **Left**: Representative whole-cell responses to 10 mM glutamate (2 seconds, –60 mV) from HEK293T cells transfected with GluA1i_γ8 tandem in control condition (black) and in presence of isatin 10 µM (red). **Right**: Paired bar plots showing effect of 10 µM isatin on desensitization τ, equilibrium current and resensitization for GluA1i_γ8. Each point is a measure of parameter in absence or presence of isatin. Bar height represents the mean value. Two-tailed paired t-test *p* values for all conditions were $p > 0.05$. **b**, Fluctuation in distances between V176 and G209 Cα residues during simulations, as a measure of stability of the binding site flanked by these residues. Data shown for both TARPg8 subunits (chains I and J). **c**, TARP γ8 contact points along its binding site, the $M4_{GluA2}$ and $M1_{GluA1}$ helices. Contacted residues are coloured depending on the number of atoms contributing to the interaction (red: high; blue: low). Contacts were computed using 'findNeighbors' in ProDy' with a 4.5 Å cutoff between heavy atoms (Bakan et al., 2011). **d**, Contact-difference maps (see Methods) for GluA1-TARP (left panel) and GluA2-TARP (right panel) from simulations. Positive values (blue) indicate contacts that are longer lived in apo vs. ligand-bound states, negative values (red) show contacts that are more persistent in ligand-bound systems compared to apo. Dotted ovals highlight the main changes in interfacial contacts from apo to ligand-bound states: ligand binding reduced TARP contact with GluA1 M1 in the top half (blue contacts in oval), but increased contact near the helix base (red contacts in oval). For GluA2 M4, ligand binding induces an overall increase in contact with TARP (red contacts in oval).

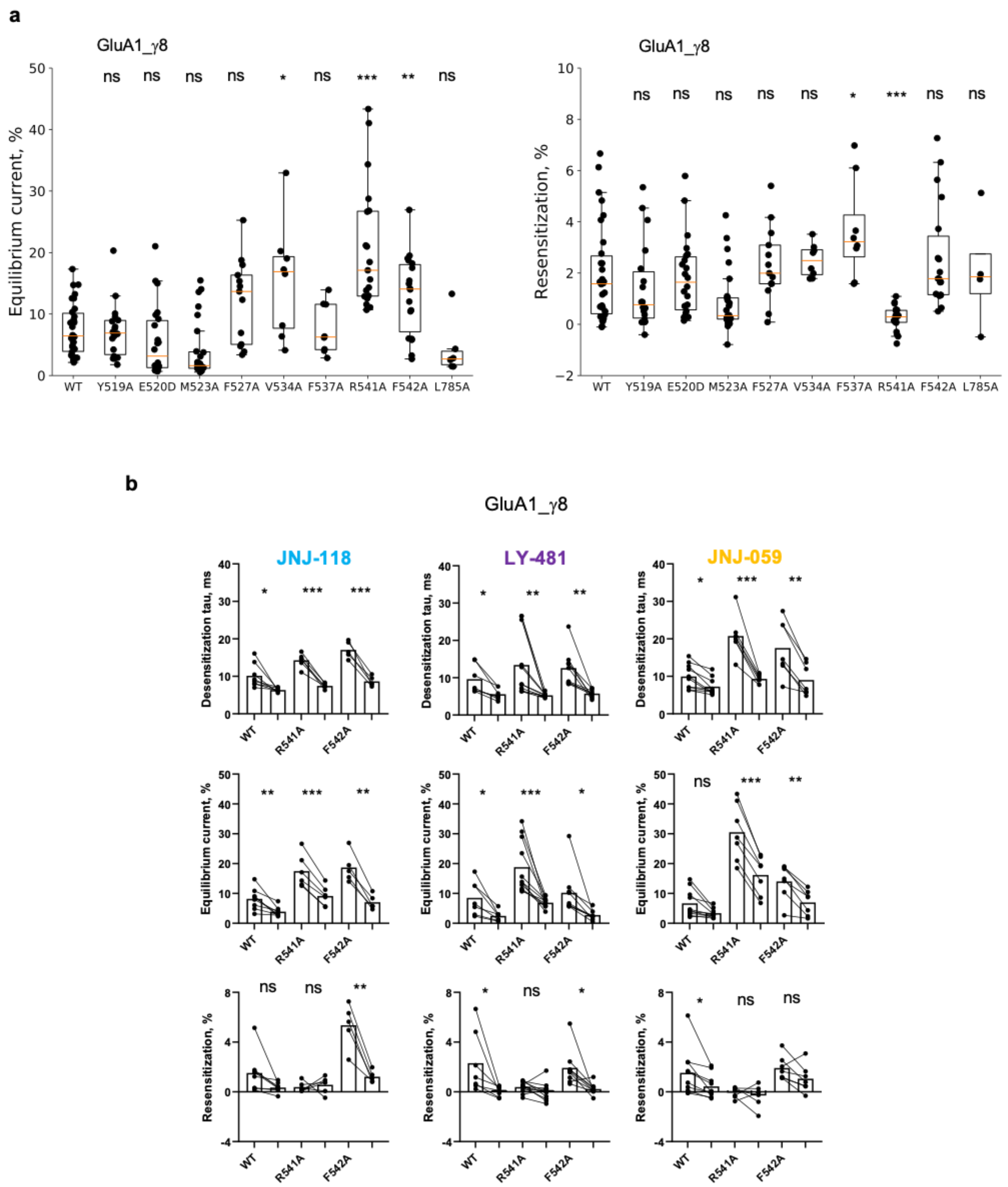


**Fig. S8. Electrophysiological analysis of GluA1 mutants in response to the three NAMs.**

**a**, and **b**, Box plots showing equilibrium current **a**, and resensitization **b**, for wild type GluA1i_γ8 and several GluA1i mutants as indicated. Boxes show the 25th/75th percentiles and whiskers indicate the furthest points that fall within 1.5 times of interquartile range from the 25th/75th percentiles. The horizontal line in each box shows the median value. Asterisks summarize one-way ANOVA test with Dunnett correction was used for multiple comparisons to wild type receptor (* *p* <= 0.05, ** *p* <= 0.01, ***

*p* <= 0.001 and ‘ns’ for p>0.05). **c**, Paired bar plots showing effect of 10 µM JNJ-118, LY-481 or JNJ-059 on desensitization $\tau$, equilibrium current and resensitization for wild type or mutated (R541A or F542A) GluA1i_γ8. Each point is a measure of parameter in absence or presence of modulator. Bar height represents the mean value. Asterisks indicate summary of two-tailed paired t-test values: * *p* <= 0.05, ** *p* <= 0.01, *** *p* <= 0.001 and ‘ns’ for p>0.05.

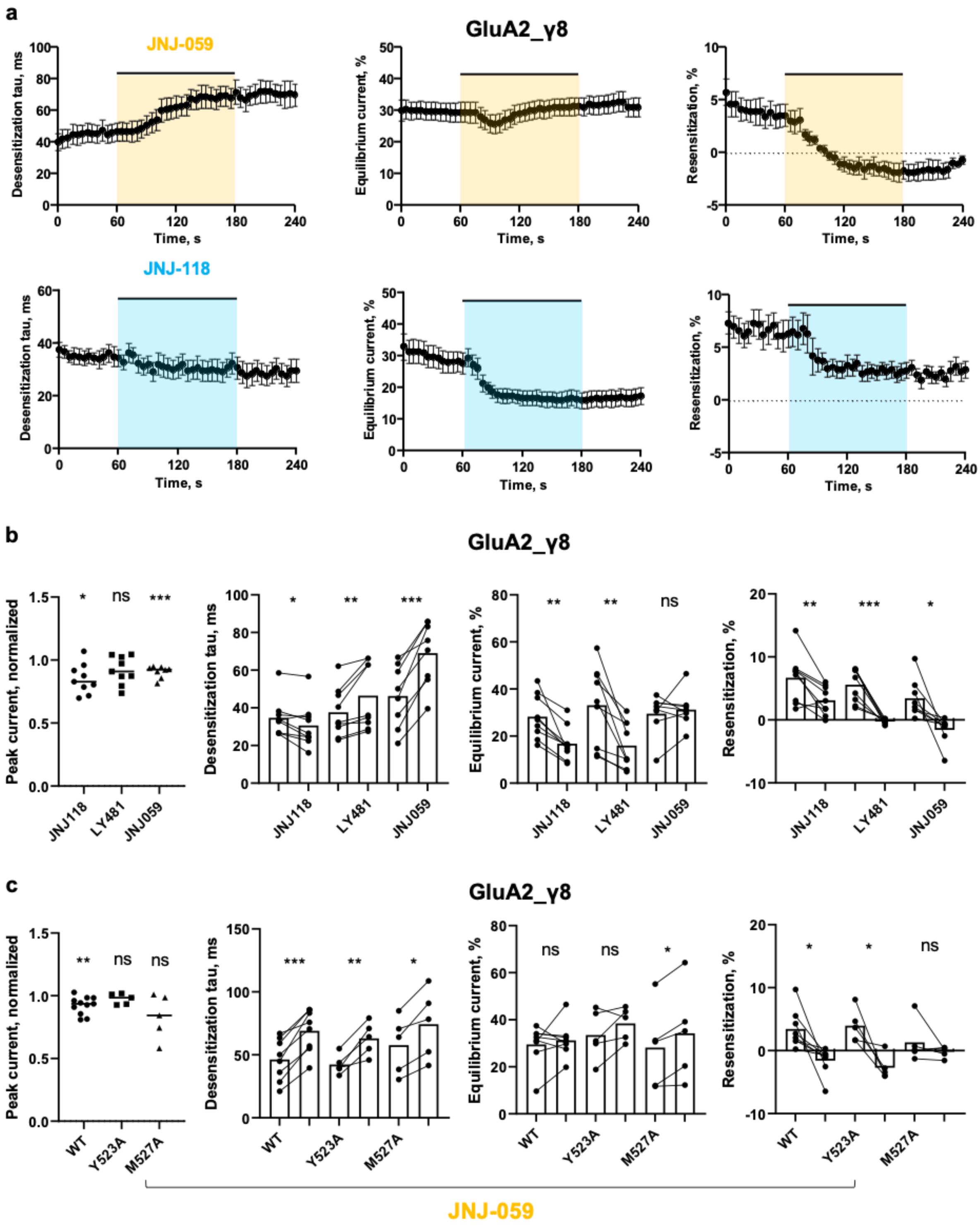


**Fig. S9. Electrophysiological analysis of GluA2 mutants in response to the three NAMs.**

**a**, Scatter plot of average desensitization $\tau$ (left), equilibrium current (middle) and resensitization (right) in time indicating the time-course of JNJ-059 (n=8; top) or JNJ-118 (n=9; bottom) effect on wild-type GluA2iQ_ γ8 receptor currents. Black circles and whiskers indicate mean values and standard error. **b**, **Left**: Pooled scatter plot of

GluA2iQ_γ8 current peak inhibition by three modulators. Each point is current peak in the presence of modulator normalized to the control peak. Horizontal lines indicate the mean values. **Right**: Paired plots showing effect of 10 µM JNJ-118, LY-481 or JNJ-059 on desensitization $\tau$, equilibrium current and resensitization for GluA2iQ_γ8. Bar height represents the mean value. **c**, **Left**: Pooled scatter plot of wild type or mutant GluA2iQ_γ8 current peak inhibition by JNJ-059. Each point is current peak in the presence of JNJ-059 normalized to the control peak. Horizontal lines indicate the mean values. **Right**: Paired plots showing effect of JNJ-059 on desensitization $\tau$, equilibrium current and resensitization for GluA2iQ_γ8. Bar height represents the mean value. Asterisks indicate summary of one-sample t-test (difference from 1; normalised peak) or two-tailed paired t-test values (paired plots): * $p <= 0.05$, ** $p <= 0.01$, *** $p <= 0.001$ and 'ns' for p>0.05.

**Table S1. Cryo-EM data collection, refinement and validation statistics.**

| | A1/2_γ8 + LY-481 Resting state | A1/2_γ8 + JNJ-118 Resting state | A1/2_γ8 + JNJ-059 Resting state | A1/2_γ8 + JNJ-059 Open state |
|---|---|---|---|---|
| | LBD-TMD (EMDB-XXX) (PDB XXX) | LBD-TMD (EMDB-XXX) (PDB XXX) | LBD-TMD (EMDB-XXX) (PDB XXX) | LBD-TMD (EMDB-XXX) (PDB XXX) |
| **Data collection and processing** | | | | |
| Microscope | FEI Titan Krios | FEI Titan Krios | FEI Titan Krios | FEI Titan Krios |
| Detector | K3 + GIF | K3 + GIF | K3 + GIF | K3 + GIF |
| Magnification | 81000X | 81000X | 81000X | 81000X |
| Voltage (kV) | 300 | 300 | 300 | 300 |
| Electron exposure (e-/Å2) | 50 | 50 | 50 | 50 |
| Defocus range (μm) | -1.2 to -2.8 | -1.2 to -2.8 | -1.2 to -2.8 | -1.2 to -2.8 |
| Pixel size (Å/pixel) | 1.07 | 1.07 | 1.07 | 1.07 |
| Symmetry imposed | C2 | C2 | C2 | C2 |
| Micrographs | 5822 | 4019 | 3781 | 13612 |
| Map resolution (Å) | 2.8 | 3.3 | 3.2 | 3.3 |
| FSC threshold | 0.143 | 0.143 | 0.143 | 0.143 |
| **Refinement** | | | | |
| Initial model used (PDB) | 7OCD | 7OCD | 7OCD | 7OCD |
| Model resolution (Å) | 2.8 | 3.3 | 3.2 | 3.3 |
| FSC threshold | 0.5 | 0.5 | 0.5 | 0.5 |
| Map sharpening B factor ($Å^2$) | -90 | -104 | -88 | -106 |
| Model composition | | | | |
| Non-hydrogen atoms | 15748 | 15962 | 15702 | 14512 |
| Protein residues | 1998 | 2008 | 1998 | 1962 |
| Ligands | LY-481: 2, ZK: 4 | JNJ-118: 2, ZK: 4 | JNJ-059: 2, ZK: 4 | JNJ-059: 2, CTZ: 4 |
| Lipids | 22 | 22 | 18 | 16 |
| B factors (Å2) | | | | |
| Protein | 45.86 | 34.91 | 52.15 | 50.23 |
| Ligand | 23.32 | 17.75 | 33.71 | 36.84 |
| R.m.s. deviations | | | | |
| Bond lengths (Å) | 0.008 | 0.006 | 0.007 | 0.007 |
| Bond angles | 0.646 | 0.796 | 0.608 | 0.583 |
| Validation | | | | |
| Molprobity score | 1.56 | 1.01 | 1.61 | 1.40 |
| Clashscore | 5.41 | 1.59 | 5.82 | 6.56 |
| Poor rotamers (%) | 0 | 0.13 | 0 | 0 |
| Ramachandran plot | | | | |
| Favoured (%) | 96.08 | 97.47 | 95.83 | 97.81 |
| Allowed (%) | 3.92 | 2.53 | 4.17 | 2.19 |
| Disallowed (%) | 0 | 0 | 0 | 0 |

**Supplementary Movie 1. Cryo-EM map of the LY-481 resting state GluA1/2_γ8 complex, outlining the NAM binding pocket (side and front views).** Colour code: GluA1-blue, GluA2-red, TARP γ8-green, LY-481-purple, lipids-grey, ligand coordinating side chains are shown in stick.

**Supplementary Movie 2. All-atom MD simulation of the LY-481 resting state GluA1/2_γ8 complex.** Colour code: GluA1-blue, GluA2-red, TARP γ8-green, LY-481-yellow (spheres).
